# Genome-Wide Transcription Response of *Staphylococcus epidermidis* to Heat Shock and Medically Relevant Glucose Levels

**DOI:** 10.1101/2024.03.18.585582

**Authors:** Kaisha N Benjamin, Aditi Goyal, Ramesh Nair, Drew Endy

## Abstract

Skin serves as both barrier and interface between body and environment. Skin microbes are intermediaries evolved to respond, transduce, or act in response to changing environmental or physiological conditions. Here, we quantify genome-wide changes in gene expression levels for one abundant skin commensal, *Staphylococcus epidermidis*, in response to an internal physiological signal, glucose levels, and an external environmental signal, temperature. We find 85 of 2354 genes change up to **∼**34-fold in response to medically relevant changes in glucose concentration (0 mM to 17 mM; adj *P* value ≤ 0.05). We observed carbon catabolite repression in response to a range of glucose spikes, as well as upregulation of genes involved in glucose utilization in response to persistent glucose. We observed 366 differentially expressed genes in response to a physiologically relevant change in temperature (37°C to 45°C; adj *P* value ≤ 0.05) and an *S. epidermidis* heat-shock response that mostly resembles the heat-shock response of related staphylococcal species. DNA motif analysis also revealed CtsR and CIRCE operator sequences arranged in tandem upstream of *dnaK* and *groESL* operons. We further identified 38 glucose-responsive genes as candidate ON or OFF switches for use in controlling synthetic genetic systems. Such systems might be used to instrument the in-situ skin microbiome or help control microbes bioengineered to serve as embedded diagnostics, monitoring, or treatment platforms.

## INTRODUCTION

Skin serves as both a barrier to the external environment and home to diverse microbial communities. Skin bacteria play a significant role in promoting and maintaining human health, contributing to skin barrier homeostasis (Zheng et al., 2022), influencing our immune system (Leech et al., 2019), and limiting pathogen invasion (Nakatsuji et al., 2017; Williams et al., 2019). One abundant skin commensal is *Staphylococcus epidermidis,* a gram-positive coagulase-negative bacterium.

*S. epidermidis* has emerged as a promising microbial chassis to enable development of engineered microbes with enhanced functionality. For example, Chen et al. engineered an *S. epidermidis* strain to produce tumor-associated antigens unique to melanoma, an aggressive type of metastatic skin cancer. When mice were colonized with the engineered *S. epidermidis* strain, a robust antitumor T cell response against localized and metastatic melanoma was generated (Chen et al., 2023). As a second example, Azitra, Inc. indicates they are engineering *S. epidermidis* strains to deliver therapeutic proteins to treat skin diseases including Netherton Syndrome and to improve skin appearance (Azitra, 2023).

Unfortunately, the tools and knowledge needed to study and reprogram *S. epidermidis* are quite limited compared to those available for established model organisms such as *Escherichia coli* or *Saccharomyces cerevisiae.* Introduction of new genes and predictable control of heterologous gene expression remain considerable challenges in bioengineering *S. epidermidis.* The nascent *S. epidermidis* knowledge base and toolkit contains methods for transformation (Monk et al., 2012; Costa et al., 2017), methods for conjugation (Brophy et al., 2018), and a small number of functionally validated promoters for control of gene expression: sarA-P1 (Bayer, Heinrichs and Cheung, 1996), P_pen_ (Meredith, Swoboda and Walker, 2008), IPTG-inducible P_spank_ (Rokop, Auchtung and Grossman, 2004), and xylose-inducible P_xylR_ (Franke et al., 2007). While successful attempts have been made to identify and characterize constitutive promoters in related staphylococcal species including *Staphylococcus aureus* (Liu et al., 2022), native transcription control elements that can serve as starting points for endogenous and dynamic control of bioengineered circuits have not yet been well characterized in *S. epidermidis*.

One application of bioengineered skin microbes could be to detect or respond to blood glucose levels, which could help in the diagnosis or treatment of diabetes. Commensal skin microbes such as *S. epidermidis* reside in subepidermal compartments of the skin with proximity to blood vessels, such as the dermis and subcutaneous adipose tissue (Nakatsuji et al., 2013; Bay et al., 2020). Such proximity could potentially facilitate the development of an engineered *S. epidermidis* strain that can sense and respond to elevated blood glucose levels (i.e., > 7mM) as a therapeutic strategy for diabetes, a chronic endocrine disorder characterized by elevated blood glucose levels and poor glycemic control (World Health Organization, 2023). To make such work practical, one would need to implement within *S. epidermidis* a transcription-based biosensor responsive to elevated blood sugar levels that results in well-regulated and rapid production of single-chain insulin. Such a use case supports the need for better characterization of glucose-inducible *S. epidermidis* regulatory elements.

Another class of applications for bioengineered skin microbes could be in response to environmental or physiological (e.g., exercise-induced) changes in temperature. With globally increasing intensity, frequency, and duration of heat waves (Perkins-Kirkpatrick and Lewis, 2020), there may be value in better understanding how commensal skin bacteria, including *S. epidermidis*, adapt and respond to increases in temperature. While the heat-shock response has been well characterized in related staphylococcal species and other prokaryotes, only three efforts have investigated the *S. epidermidis* heat-shock response by using semi-quantitative protein assays (Ooronfleh, Streips, and Wilkinson, 1990), focusing on only a small number of genes (Vandecasteele et al., 2001) or using comparative genomics (Chastanet, Fert, and Msadek, 2003). We thus chose to also quantitatively explore the genome-wide transcription response of *S. epidermidis* to heat shock, both as a reference case for glucose response and, for its own merits.

We investigated the genome-wide transcription response in the non-biofilm forming, nonpathogenic *S. epidermidis* strain (ATCC 12228) to heat shock and medically relevant glucose concentrations. We performed RNA sequencing on samples exposed to a sudden temperature increase and a glucose challenge to investigate the ability of the organism to adapt and respond to changing environmental conditions. We used differential expression analysis of samples taken during the mid-exponential growth phase to identify candidate genes that are either upregulated or downregulated in response to each condition. We further curated a subset of glucose-responsive genes that might serve as templates for ON or OFF switches.

## MATERIALS AND METHODS

### Bacterial Strain and Culture

We started each *S. epidermidis* ATCC 12228 culture from a fresh colony plate (< 7 days old) using a single colony. We used Tryptic Soy Broth (TSB) without Dextrose (BD 286220) as the culture medium for all experiments.

### Heat-Shock Experiments

We grew overnight broth cultures in fresh medium supplemented with 0.2% w/v glucose for 18 h at 37°C with shaking. Cultures were then diluted 32-fold in fresh medium supplemented with 0.2% w/v glucose and grown at 37°C with shaking. When cultures were in mid-exponential phase (OD_600_ ∼ 0.5), we transferred them to pre-warmed Erlenmeyer flasks followed by incubation at 45°C for 10 minutes. We then harvested cultures for RNA sequencing (below). Control cultures in mid-exponential phase were not exposed to heat shock but instead were immediately harvested for RNA sequencing. We performed our heat-shock experiments in triplicate to generate three biological replicates.

### Glucose Challenge Experiments

We grew overnight broth cultures in fresh medium supplemented with 13.9 mM glycerol for 25 h at 37°C with shaking. We then diluted cultures 50-fold in fresh medium supplemented with 13.9 mM glycerol and continued growth at 37°C with shaking. When cultures were in mid-exponential phase (OD_600_ ∼ 0.5), we added glucose and measured the glucose concentration (2 mM, 5 mM, 10 mM, 17 mM, or 50 mM) of each culture using the Contour NEXT ONE Blood Glucose Monitoring System. We added an equivalent volume of fresh medium lacking glucose to the control cultures. We grew cultures at 37°C with shaking for an additional 20 minutes and then harvested for RNA sequencing (below). We performed our glucose challenge experiments in triplicate to generate three biological replicates.

### Step-down Experiments

We grew overnight broth cultures in fresh medium supplemented with 13.9 mM glycerol for 25 h at 37°C with shaking. We diluted cultures 50-fold in fresh medium supplemented with 13.9 mM glycerol and continued growth at 37°C with shaking. When cultures were in mid-exponential phase (OD_600_ ∼ 0.5), we added glucose and measured the glucose concentration (10 mM) of each culture using the Contour NEXT ONE Blood Glucose Monitoring System. Cultures were then grown at 37°C with shaking for 20 minutes and then pelleted at 5,000 xg for 10 minutes at 24°C. We then resuspended the pellets in fresh medium supplemented with 2 mM glucose. We grew cultures at 37°C with shaking for an additional 20 minutes and harvested for RNA sequencing (below). We used the 10 mM glucose challenge condition (above) as the control condition for our step-down experiments. We performed our step-down experiments in triplicate to generate three biological replicates.

### Batch Culture Experiments

We grew overnight broth cultures in fresh medium supplemented with glucose (0.2% w/v or 1% w/v) for 18 h at 37°C with shaking. We measured the glucose concentration of each culture using the Contour NEXT ONE Blood Glucose Monitoring System. We diluted cultures 32-fold in fresh medium supplemented with glucose (0.2% w/v or 1% w/v) and grew at 37°C with shaking. We harvested mid-exponential phase cultures (OD_600_ ∼ 0.5) for RNA sequencing (below). We performed our batch culture experiments in duplicate to generate two biological replicates.

### RNA Stabilization and Extraction

Immediately after each experiment, we pelleted samples by centrifugation at 5,000 x g for 10 minutes at 4°C and then resuspended the pellets in RNAlater (Invitrogen AM7021); samples were incubated in RNAlater at 4°C for 24 h. After incubation, we pelleted samples by centrifugation at 5,000 x g for 10 minutes at 4°C and resuspended the pellets in 1µl of 100X TE Buffer, 50 μl of lysostaphin (1 mg ml^-1^), and 50 μl of mutanolysin (5KU ml^-1^). We performed lysis for 25 minutes at 37°C with vortexing at 5-minute intervals. We then treated samples with 25 µl of Proteinase K (Qiagen 19131) and incubated for an additional 30 minutes at 37°C. We added 700 μl of Buffer RLT (Qiagen 79216) to each sample and vortexed vigorously for 5 to 10 seconds. We transferred the resulting suspension to a 2 ml Safe-Lock tube (Eppendorf 0030123620) and mechanically disrupted the samples using a TissueLyser LT (Qiagen 85600) for 5 minutes at maximum speed with intervals of 30 seconds of bead beating and 30 seconds of resting on ice. After bead beating, we centrifuged the samples in an Eppendorf MiniSpin (022620100) for 15 seconds at maximum speed (12,100 x g) and then transferred the supernatant to a new tube. We mixed the supernatant well with an equal volume of 100% ethanol by pipetting. We applied this mixture to a RNeasy Mini spin column and extracted RNA according to the manufacturer’s instructions using a RNeasy Mini Kit (Qiagen 74106). We performed on-column DNase digestion using the RNase-Free DNase Set (Qiagen 79254). We eluted samples in RNase-free water according to the manufacturer’s instructions and stored recovered RNA at −80°C until library preparation. We used RNaseZap RNase Decontamination Solution (Invitrogen AM9780) on all surfaces to prevent RNA degradation. RNA quality was analyzed using an Agilent Bioanalyzer and quantified by a Qubit fluorometer according to manufacturer’s instructions. Our RNA integrity number (RIN) values ranged from 8.0 to 10.

### Library Preparation and Sequencing

We used Novogene Co., LTD (Beijing, China) to carry out our rRNA depletion, cDNA library preparation, and sequencing as part of their Prokaryotic RNA Sequencing service. cDNA libraries were sequenced on an Illumina NovaSeq 6000 Sequencing System with a 150 bp paired-end run configuration to a depth of ∼30 million reads.

### Raw Sequence Data Quality Control & Processing

We processed raw reads (FASTQ files) using FastQC v0.12.1 (Andrews, 2010) with default settings to assess initial read quality and then examined the results using MultiQC v1.14 (Ewels et al., 2016). We processed FASTQ files using Trim Galore v0.6.10 (Krueger, 2012) with default settings to trim low-quality (Phred score < 20) ends from reads and to trim auto-detected adapters. Reads that became shorter than 20 bp because of either quality or adapter trimming were discarded.

### Reference Genome for Mapping

We used the *Staphylococcus epidermidis* ATCC 12228 genome assembly ASM987345v1 (GenBank accession GCA_009873455.1, RefSeq accession GCF_009873455.1) from NCBI in the FASTA format along with information on genes and other features in the GFF format. The genome consists of a chromosome (GenBank accession CP043845.1, RefSeq accession NZ_CP043845.1) of size 2,504,425 bp and a plasmid (GenBank accession CP043846.1, RefSeq accession NZ_CP043846.1) of size 21,978 bp. We converted GFF features to GTF format by using the *gffread* program in the Cufflinks v2.2.1 package (Trapnell et al., 2010) and to BED format by using the AGAT v1.0.0 toolkit (Dainat, 2019) for use in downstream analysis.

### Mapping and Transcript Quantification

We used Bowtie2 v2.5.1 (Langmead and Salzberg, 2012) to build a Bowtie index from the *S. epidermidis* ATCC 12228 genome assembly ASM987345v1 before mapping the RNA-Seq reads in the paired-end FASTQ files to this reference genome using default settings. The resulting BAM files were coordinate-sorted and indexed; alignment summary statistics were reported using SAMtools v1.17 (Danecek et al., 2021). We ran RSeQC v5.0.1 (Wang, Wang, and Li, 2012) on the sorted BAM files to determine the strandedness of the reads for the strand-specific RNA-seq data. We used *featureCounts* in the Subread v2.0.6 package (Liao, Smyth, and Shi, 2013) to count mapped reads at both the transcript and gene levels from sorted BAM files for genomic features such as CDSs, based on previously determined read strandedness. We merged counts from each sample at both the transcript and gene levels. We used the resulting merged count matrices in subsequent differential expression analysis.

### BLASTP Homology Search

The KEGG Pathway Database (Kanehisa and Goto, 2000) Genome Entry T00110 (Org code: sep) lists genome assembly ASM764v1 (GenBank accession GCA_000007645.1, RefSeq accession GCF_000007645.1) as the reference genome for *S. epidermidis* ATCC 12228. Genome assembly ASM764v1 uses alternate gene designations compared to the genome assembly ASM987345v1 used in this study. To leverage KEGG pathway gene sets for Gene Set Enrichment Analysis (GSEA), we conducted a BLASTP homology search between the two genome assemblies using NCBI BLAST+ executable v2.14.0+ (Camacho et al., 2009) to find genes in genome assembly ASM987345v1 with the highest degree of homology to genes in genome assembly ASM764v1 thereby enabling cross-mapping of the genes represented in KEGG Pathway Gene Sets.

### Differential Expression Analysis

We used principal component analysis (PCA) to first visualize the expression data; we applied a regularized log (rlog) transformation to all expression data. We then visualized sample-to-sample distances via PCA and found that one replicate from the step-down experimental condition was over 4-fold off on the second principal component against all other experimental samples, and over 10-fold off on the first principal component against the other two step-down samples (Figure S1). We thus excluded the data from this one step-down replicate in all further analyses. We then analyzed data from non-transformed count matrices using the DESeq2 R package (Love, Huber, and Anders, 2014), which can evaluate differential expression on as few as two biological replicates. We defined differentially expressed genes (DEGs) of significance using the following criteria: |log2 fold change| (i.e., log2FC) ≥ 1.5 and adjusted *P* value ≤ 0.05. We applied the apeglm (log fold change shrinkage) method (Zhu, Ibrahim, and Love, 2018) to the raw counts to stabilize variability in log fold change calculations. We then constructed volcano plots using the EnhancedVolcano R package (Blighe, Rana, and Lewis 2023) and further customized them using ggplot2 (Wickham, 2016). We designed Circle plots using shinyCircos (Yu, Ouyang, and Yao, 2017). We also constructed the two scatter plots, visualizing the relationship between the heat-shock and G17 experimental conditions and between the step-down and G2 experimental conditions, using ggplot2.

### Pathway and Gene Identification

We explored gene functions using the KEGG and GO pathways database and manually curated a gene annotation table, drawing from the KEGG (organism code *sep*), BioCyc (GCF_000007645), and UniProt databases. After determining gene-to-pathway annotations, we used the GSEA tool (Subramanian et al., 2005; Mootha et al., 2003) and the fgsea R package (Korotkevich et al., 2021) to conduct gene set enrichment analysis. We used Fisher’s method to combine results that overlapped across GSEA and fgsea, creating a single *P* value that reflected the two independent adjusted *P* values. We reduced GO term redundancy using REVIGO (Supek et al., 2011), with default parameters and a “small (0.5)” resulting list. Once KEGG and GO enriched pathways were identified, we performed independent research to cross-validate the results and combined pathways that were identified in both KEGG and GO databases.

### Switch Identification

We identified switches using the DRomics package, a tool used for concentration-response (or dose-response) characterization from -omics data (Marie Laure Delignette-Muller et al., 2023; Floriane Larras et al., 2018). We modeled all genes with an absolute log fold change ≥ 2. We performed a rlog transform on gene counts and then used DRomics to identify the appropriate best-fit monophasic or biphasic model; genes that failed to model due to a slope near zero were deemed dose-insensitive.

### Batch Culture Bioinformatics Analysis

Novogene (Beijing, China) completed bioinformatics analyses for our batch culture experimental condition as part of their Prokaryotic RNA Sequencing standard analysis. **Raw Sequence Data Quality Control:** Novogene processed raw reads (FASTQ files) using Fastp (Chen et al., 2018). Clean data for downstream analysis were obtained by removal of low-quality reads, adapters, and poly-N sequences. **Reference Genome and Mapping:** Novogene obtained the reference genome (GenBank accession GCA_009873455.1, RefSeq accession GCF_009873455.1) and gene model annotation files from NCBI and aligned clean reads to the reference genome using Bowtie2 (Langmead and Salzberg, 2012). **Transcript Quantification:** Novogene used *FeatureCounts* (Liao, Smyth, and Shi, 2013) to count reads mapped to each gene and then calculated the fragments per kilobase of transcript per million fragments mapped (*FPKM*) of each gene based on gene length and read counts mapped to the gene (Trapnell et al., 2010). **Differential Expression Analysis:** Novogene performed differential expression analysis using the DeSeq2 R package (Love, Huber, and Anders, 2014) and adjusted *P* values using the Benjamini and Hochberg method for controlling the false discovery rate (Benjamini and Hochberg, 1995). Differentially expressed genes (DEGs) of significance were defined using the following criteria: |log2 fold change| (i.e., log2FC) ≥ 1.5 and adjusted *P* value < 0.05.

### Data Deposition and Availability

The original contributions presented in the study are publicly available. The data discussed in this publication have been deposited in NCBI’s Gene Expression Omnibus (Benjamin et al., 2024) and are accessible through the GEO Series accession number GSE261664.

## RESULTS

The heat-shock response (HSR), a transcription program observed in several eukaryotes and prokaryotes, is a crucial strategy whereby cells adapt to a sudden temperature increase or other environmental stresses (Cao et al., 1999). The HSR helps cells maintain protein homeostasis by protection from heat-induced protein denaturation, misfolding, and aggregation. HSR has been studied in detail in *Escherichia coli*, *Streptomyces* spp., and *Bacillus subtilis* (Lemaux et al., 1978; Guglielmi et al., 1991; Schumann, 2003). While the HSR is highly conserved across prokaryotes, the regulatory mechanisms that govern the expression of heat-shock genes exhibit great diversity among bacterial species (Roncarati and Scarlato, 2017; Schumann, 2016). Prior studies of the HSR in *S. aureus* (Chastanet, Fert, and Msadek, 2003; Anderson et al., 2006; Fleury et al., 2009) and the gram-positive model organism *B. subtilis* provide a context from which to increase our understanding of the HSR of *S. epidermidis* and other low-GC content gram-positive bacteria.

### Differential Gene Expression in *S. epidermidis* Under Heat Stress

To identify differentially expressed genes in heat-shocked *S. epidermidis* ATCC 12228 cells, we shifted mid-exponential phase cells from physiological growth (37°C) to heat-shock conditions (45°C) for 10 minutes (Figure 1A). We used RNA sequencing to analyze gene expression profiles and then compared the expression profiles of heat-shocked cells to those of unstressed cells. Differentially-expressed genes (DEGs) of significance were defined using the following criteria: |log2 fold change| (i.e., log2FC) ≥ 1.5 and adjusted *P* value ≤ 0.05. By these criteria, we identified 366 of 2354 genes (∼15.5% of the genome) with log2FC values ≥ 1.5, among which 235 were upregulated and 131 were downregulated (Table S1, Table S2). Downregulated and upregulated genes were expressed over a −4 to +6 log2FC range (Figure 2A).

**Figure 1.**
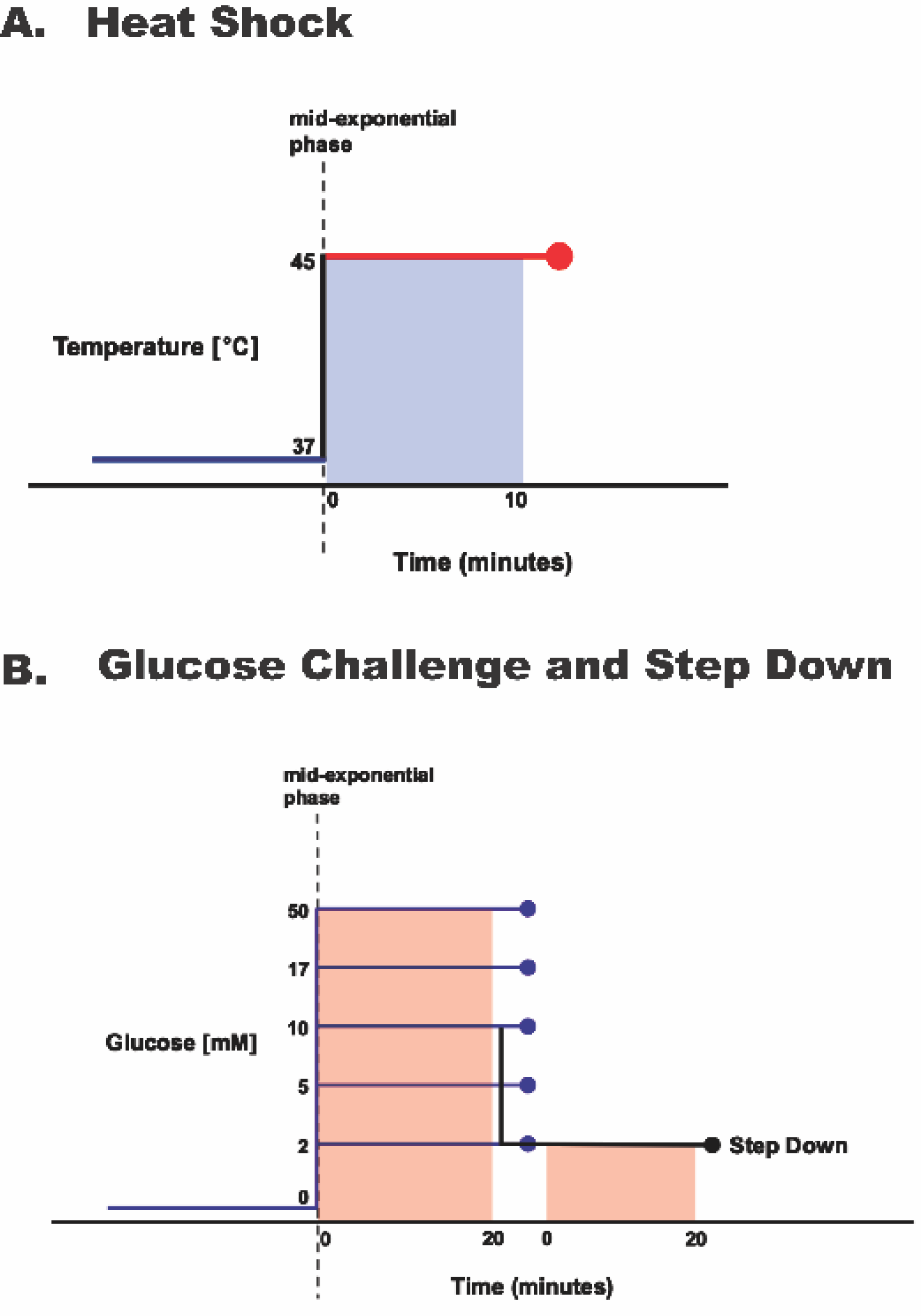
Environmental Perturbation of *Staphylococcus epidermidis*. Log-phase cultures were exposed to **(A)** a 10-minute increase in temperature from 37°C to 45°C or **(B)** a range of 20-minute glucose spikes (concentrations as noted) and a 10 mM spike followed by a step down to 2 mM.

**Figure 2.**
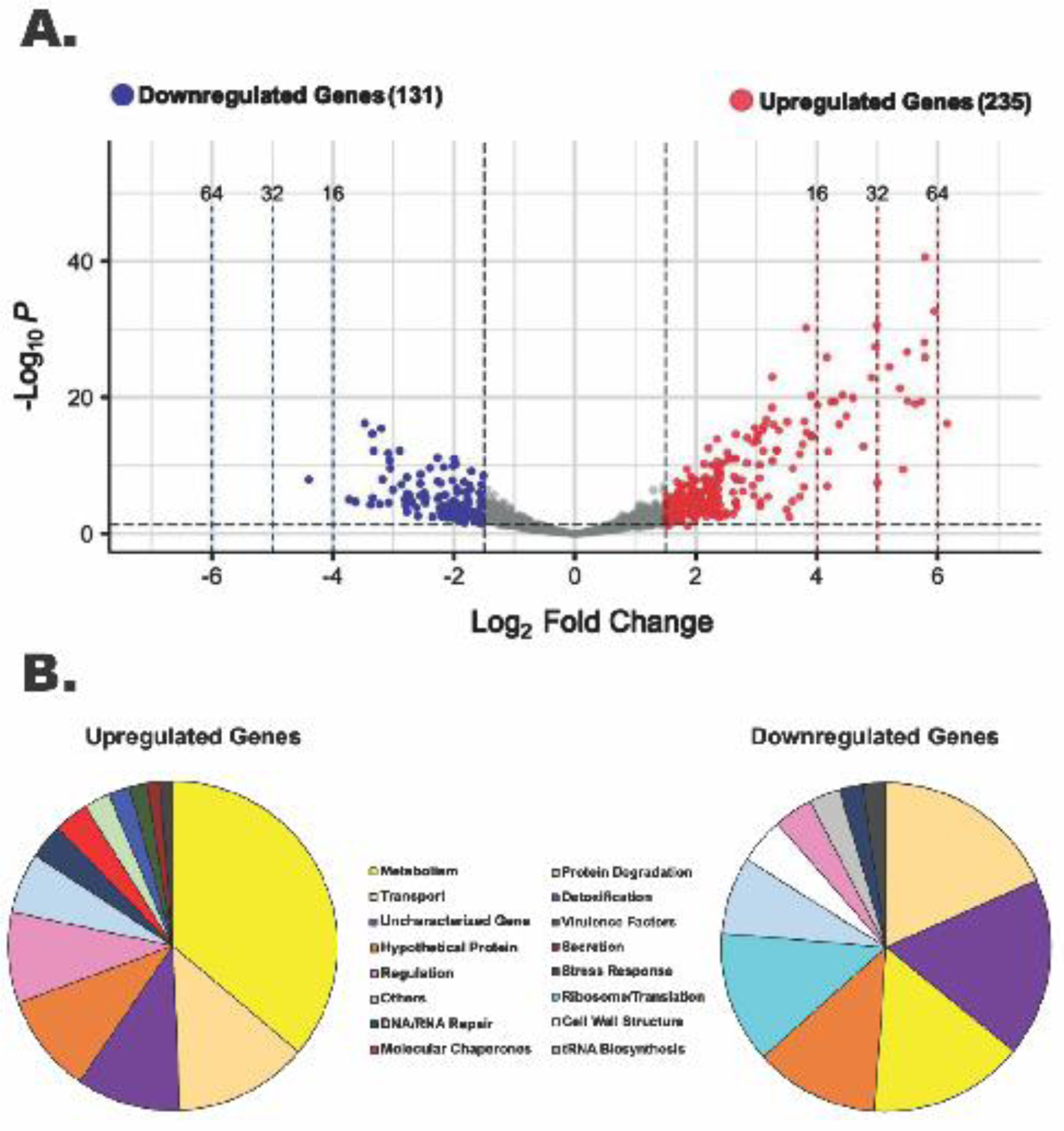
A Sudden Temperature Increase Causes Transcript Levels to Change up to ∼71-fold. **(A)** Volcano plot showing the differentially expressed genes (DEGs) for the heat-shock experimental condition with |log2 FC|≥ 1.5 and adjusted *P* value ≤ 0.05 as the threshold. The red dots represent 235 significantly upregulated genes, and the blue dots represent 131 significantly downregulated genes**. (B)** Summary of the significantly upregulated and downregulated genes during the heat-shock response in *S. epidermidis* assigned to functional groups according to GO and KEGG pathways (in %).

We observed increased expression of several heat-shock genes well-characterized in other organisms (Anderson et al., 2006; Fleury et al., 2009; Schumann, 2003). For example, transcript levels of the *dnaK* (*hrcA*, *grpE*, *dnaK*, *dnaJ*, *prmA*), *groESL* (*groES*, *groL*), and *clpC* (F1613_RS04215 (*CtsR* family transcription regulator), F1613_RS04220 (UvrB/UvrC motif-containing protein), F1613_RS04225 (protein arginine kinase), F1613_RS04230 (ATP-dependent Clp protease ATP-binding subunit *clpC*)) operons, encoding the major cell chaperones and proteases, were upregulated ∼8-15, ∼10-11, and ∼42-53 absolute fold, respectively (Table S1). Other known heat-shock genes including *clpB*, *clpP*, the Hsp33 family molecular chaperone *hslO*, and MecA, an adaptor protein necessary for ClpC chaperone activity (Schlothauer et al., 2003) were upregulated by 71-, 8.9-, 4.14-, and 3.84-fold, respectively (Table S1). Among the most upregulated genes (**∼**22-61-fold) were members of the *lac* operon (*lacA, lacB,* F1613_RS11920 (tagatose-6-phosphate kinase), *lacD*, F1613_RS11910 (PTS lactose/cellobiose transporter subunit IIA), F1613_RS11905 (lactose-specific PTS transporter subunit EIIC), *lacG*), vraX, F1613_RS03870 (ArgE/DapE family deacylase), cytochrome ubiquinol oxidase subunits I and II (F1613_RS06745 and F1613_RS06750), F1613_RS01555 (MarR family transcription regulator), F1613_RS12445 (hypothetical protein), F1613_RS01550 (NAD(P)/FAD-dependent oxidoreductase), and F1613_RS03780 (MFS transporter) (Table S1).

We observed other upregulated genes of potential interest. For example, *BlaZ*, *blaI*, and *blaR1*, components of the *bla* operon that encode for a β-lactamase (Llarrull, Prorok and Mobashery, 2010) were upregulated ∼4.8-18.3-fold. Members of the urease operon (F1613_RS12320, *ureE*, F1613_RS12330) along with two competence protein ComK orthologs (F1613_RS10000 and F1613_RS06475) displayed increased transcript levels, consistent with previous observations of genes induced by heat shock in *S. aureus* (Anderson et al., 2006; Fleury et al., 2009). Twenty-three hypothetical proteins and 24 uncharacterized genes (47 total) were also upregulated under heat-shock conditions.

Among the most downregulated genes (**∼**10-21-fold) were F1613_RS05940 and *dltABCD*, components of the *dlt* operon required for the d-alanylation of teichoic acids in gram-positive bacterial cell walls (Kovacs et al., 2006) (Table S2). Several genes encoding ribosomal proteins (*rplJ, rplL*, *rplT*, *rpmI*, *rpsF*, *rpsO*, *rpsR*) and tRNA-ligases (*ileS*, *thrS*, *serS*) were also downregulated (**∼**2.9-8.3-fold) (Table S2), consistent with the transient inhibition of protein synthesis that occurs in response to heat shock in other organisms (Duncan and John W.B. Hershey, 1989). Components of the *psm*β operon (F1613_RS07060, F1613_RS07065, F1613_RS07070, F1613_RS07075) that encode for β-class phenol-soluble modulins (PSMs) (Cheung et al., 2014; Wang et al., 2011), and the PSM transporter system (*pmtA*, *pmtB*, and *pmtC*) (Chatterjee et al., 2013) were downregulated ∼3-5-fold. In total, 24 genes involved in transport were downregulated up to ∼11-fold (Table S2), with more than half of them belonging to the ATP-binding cassette (ABC) transporter superfamily. Two cold-shock genes (*cspA* and F1613_RS05710) displayed decreased transcript levels, consistent with previous observations of genes repressed by heat shock in *S. aureus* (Fleury et al., 2009). Two helix-turn-helix transcription regulators (F1613_RS10440 and F1613_RS09035) were downregulated ∼8.5 and ∼3.5-fold, respectively (Table S2). We also observed downregulation of other transcription regulators including *rsp*, F1613_RS11065 (GntR family transcription regulator), and *pyrR* by 5.5-, 4.6-, and 4.1-fold respectively (Table S2). Sixteen hypothetical proteins and 23 uncharacterized genes (39 total) were also downregulated under heat-shock conditions.

### Functional Classification of Differentially Expressed Genes in *S. epidermidis* Under Heat Stress

The genome of *Staphylococcus epidermidis* ATCC 12228 contains 2354 protein-coding genes, of which 207 are hypothetical and 71 are uncharacterized (278 total or ∼12% of all genes), indicating their biological functions are unknown or not yet established. We manually grouped 280 of 366 heat shock DEGs (∼77%) into functional groups using GO and KEGG databases (Figure 2B); 23% of heat shock DEGs had no assigned functions. We observed known functional classes that are upregulated under heat-shock conditions in all domains of life (Richter, Haslbeck, and Buchner, 2010), namely Metabolism, Transport, Regulation, DNA/RNA Repair, Molecular Chaperones, Protein Degradation, and Detoxification (Figure 2B). A significant proportion (85; ∼36%) of upregulated genes were involved in metabolism, including sugar, amino acid, and fatty acid metabolism (Table S1; Figure S2). We also observed increased expression of genes in the Virulence Factors, Secretion, and Stress Response functional classes (Figure 2B). Ribosome/Translation, tRNA Biosynthesis, and Ribosome Biogenesis functional classes accounted for a significant proportion (22; ∼17%) of downregulated genes (Figure 2B; Table S2), consistent with a transient inhibition of protein synthesis. Genes involved in Transport, Metabolism, Cell Wall Structure, Regulation, DNA/RNA Repair, and Stress Response were also downregulated under heat-shock conditions (Figure 2B). We assigned DEGs grouped into minor functional classes that contained only a small number of genes to the “Others” category in each pie chart (Figure 2B). Fourteen upregulated genes and 10 downregulated genes were assigned to the “Others” category and their functions are detailed in the supplementary material (Table S1; Table S2).

### Transcription Responses to Glucose in *S. epidermidis*

Six-carbon sugars (hexoses) such as glucose are the preferred carbon and energy sources for many prokaryotes including *S. epidermidis*. Prior studies in staphylococcal species demonstrated that glucose utilization supports faster growth and higher metabolic rates (Halsey et al., 2017). The presence of glucose also inhibits the expression of genes required for uptake and utilization of alternative carbon sources, an adaptive regulatory mechanism called carbon catabolite repression (CCR) (Görke and Stülke, 2008). We performed RNA sequencing on cultures exposed to 20-minute glucose spikes across a range of concentrations and to persistent glucose to better understand the ability of *S. epidermidis* to adapt and respond to glucose. Our underlying goal was to support development of commensal microbes bioengineered to diagnose, monitor, or treat diabetes.

### Identifying Genes that Might be Useful Starting Points for Controlling Bioengineered Bacteria in Treating Diabetes

We challenged mid-exponential phase cells by subjecting them to 2 mM, 5 mM, 10 mM, 17 mM, or 50 mM glucose spikes for 20 minutes (Figure 1B). We used RNA sequencing to analyze gene expression profiles and compared the resulting expression profiles of glucose-challenged cells to those of unchallenged cells (Figure 3A). Differentially expressed genes (DEGs) of significance were identified using the following criteria: |log2 fold change| (i.e., log2FC) ≥ 1.5 and adjusted *P* value ≤ 0.05 (Table S3-S7). We examined rlog transformed counts data from the medically relevant (G2-G17) glucose concentrations, searching for candidate transcripts that might be potential starting points for glucose-responsive switches. We found 38 potential switches by modeling all genes with absolute log2 fold change values ≥ 2 in at least one medically relevant glucose challenge experimental condition (Figure S3).

**Figure 3.**
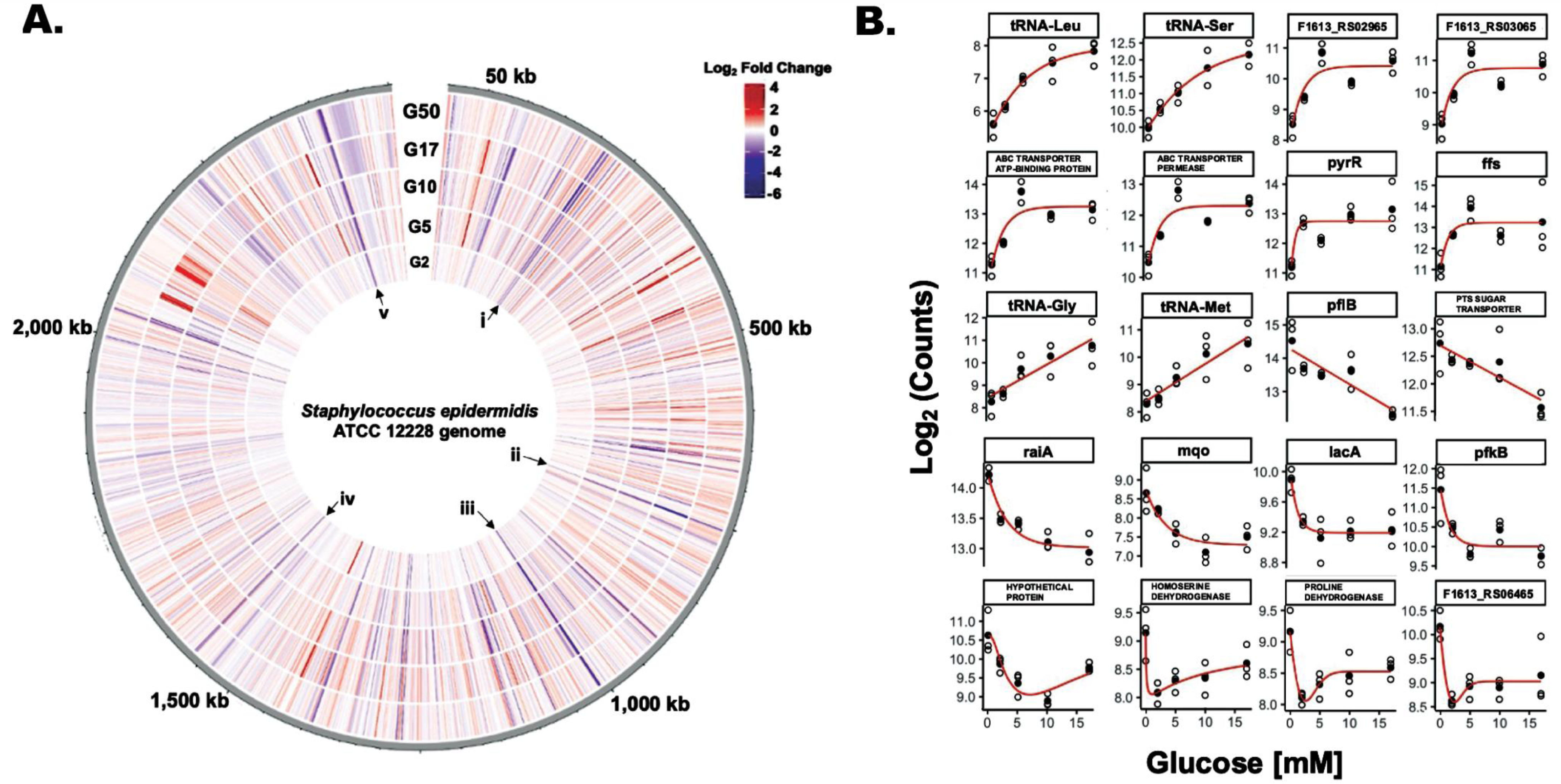
Eighty-five *S. epidermidis* Genes Change Expression Levels in Response to Glucose. **(A)** Circular transcriptome map showing normalized gene expression levels in the *S. epidermidis* genome in response to glucose. Log2 fold change relative to control for cells exposed to 2 mM (G2), 5 mM (G5), 10 mM (G10), 17 mM (G17), or 50 mM (G50) glucose spikes. Each bar denotes a single gene; red bars represent significantly upregulated genes and blue bars represent significantly downregulated genes. Roman numerals **i** (*sdaAB, rbsU*), **ii** (*pflB*), **iii** (*glpR-pfkB* operon), **iv** (F1613_RS07845 (homoserine dehydrogenase), and **v** (members of the *lac* operon) correspond to select groups of genes that are downregulated across all five glucose spike conditions. **(B)** Glucose concentration-response curves for a representative subset of genes that have potentially interesting glucose-responsive switch properties.

We selected twenty genes as representative candidates with potentially interesting glucose-responsive switch properties (Figure 3B). Among the potential switches that exhibited an OFF-to-ON transition were two DUF2871 domain-containing proteins (F1613_RS03065 and F1613_RS02965), F1613_RS00340 (ABC transporter ATP-binding protein), F1613_RS00345 (ABC transporter permease), *pyrR* (bifunctional pyr operon transcriptional regulator), *ffs (*signal recognition particle sRNA), and four tRNA genes. We also identified genes likely subject to carbon catabolite repression (CCR) that might serve as potential ON-to-OFF switches, including F1613_RS01060 (PTS sugar transporter subunit IIC), *lacA*, *pfkB*, and F1613_RS09950 (proline dehydrogenase) (Görke and Stülke, 2008; Nuxoll et al., 2012). Other promising ON-to-OFF switch candidates include *pflB* (formate C-acetyltransferase), *raiA* (ribosome-associated translation inhibitor), *mqo* (malate dehydrogenase (quinone)), F1613_RS05750 (hypothetical protein), F1613_RS07845 (homoserine dehydrogenase), and F1613_RS06465 (IDEAL domain-containing protein) (Figure 3B). We examined counts data from the medically relevant (G2-G17) glucose concentrations and also noted a class of genes whose expression did not change in response to a glucose spike compared to an unchallenged (0 mM) control. These glucose-independent genes included *lqo* (L-lactate dehydrogenase (quinone)), F1613_RS08490 (transglycosylase domain-containing protein), *typA* (translational GTPase TypA), *rnr* (ribonuclease R), and *noc* (nucleoid occlusion protein).

### Genes Repressed in Response to 20-minute Glucose Spikes

We observed 18 genes that were downregulated across all five glucose spike conditions and an additional ten genes that were downregulated across the top four glucose spike conditions (Figure 4B; Figure S4). For example, genes involved in **lactose metabolism** (F1613_RS11920 (tagatose-6-phosphate kinase), *lacB*, and *lacA*), **ribose transport** (*rbsU*, *rbsD*), **fructose utilization** (F1613_RS05160 (PTS fructose transporter subunit IIABC), *pfkB*, and F1613_RS05150 (DeoR/GlpR family DNA-binding transcription regulator)), **proline catabolism** (F1613_RS09950 (proline dehydrogenase)), **the glyoxalase pathway** (F1613_RS05685 (glyoxalase)), **the succinate dehydrogenase complex** (F1613_RS07025 (succinate dehydrogenase cytochrome b558 subunit)), and **ethanol degradation** (*adhP*) were downregulated, consistent with previous observations of gene expression changes that occur during CCR (Gutierrez-Ríos et al., 2007; Penninckx, Jaspers, and Legrain, 1983; Nam, 2005; Arndt and Eikmanns, 2007; Görke and Stülke, 2008; Nuxoll et al., 2012; Halsey et al., 2017) (Table S3-S7). We also observed decreased expression of *sdaAB* (L-serine ammonia-lyase iron-sulfur-dependent subunit beta), *raiA*, F1613_RS03360 (universal stress protein), F1613_RS00870 (GntR family transcription regulator), F1613_RS06465 (IDEAL domain-containing protein), F1613_RS10135 (AAA family ATPase), F1613_RS07845 (homoserine dehydrogenase), F1613_RS10140 (DUF4238 domain-containing protein), F1613_RS06500 (fatty acid desaturase), and genes involved in formate metabolism (*pflA* and *pflB)* across at least four glucose spike conditions. Four hypothetical proteins and one uncharacterized gene (five total) were also downregulated across at least four glucose spike conditions (Table S3-S7).

**Figure 4.**
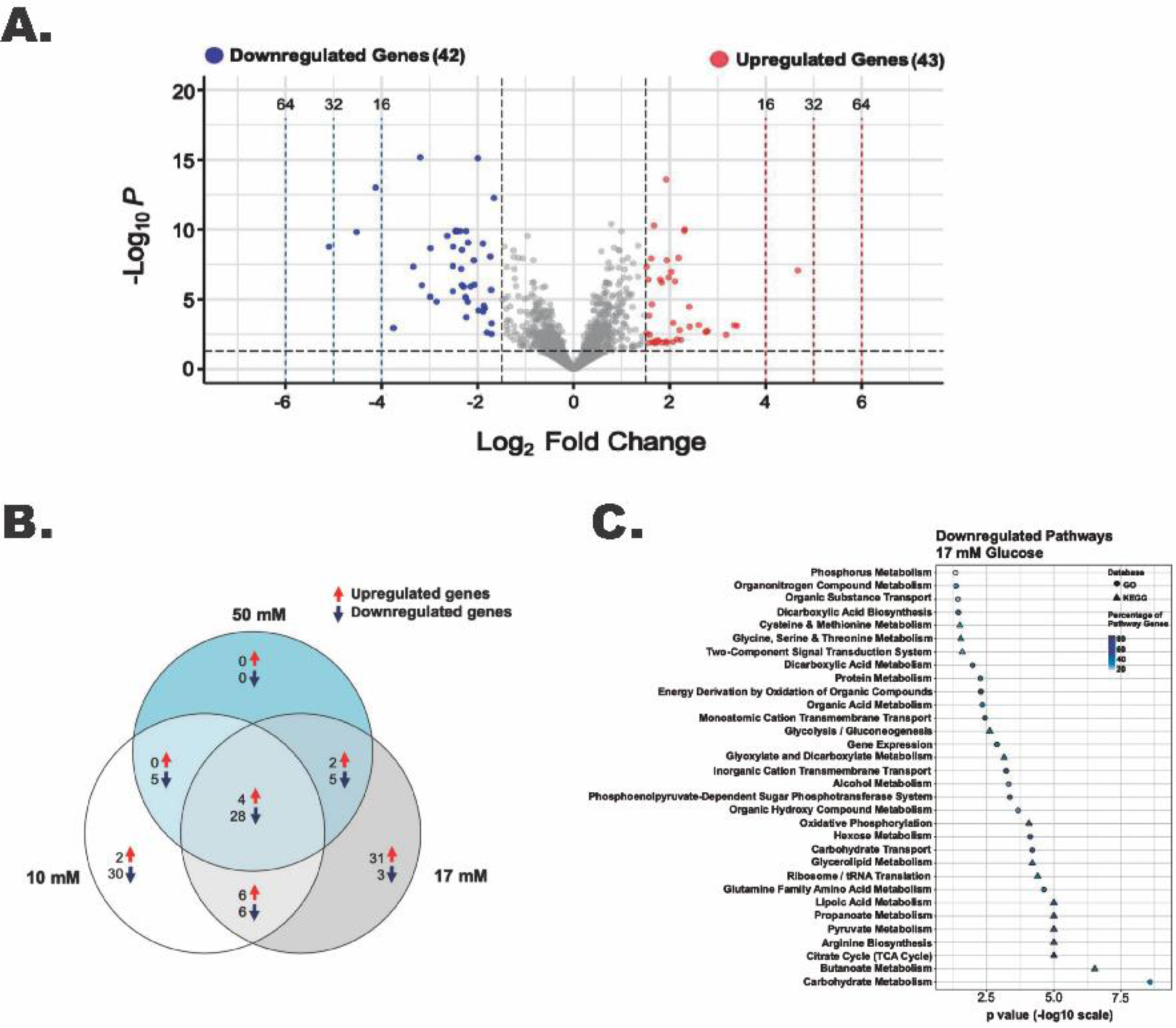
A 17 mM Glucose Spike Causes Transcript Levels to Change up to ∼34-fold. **(A)** Volcano plot showing the differentially expressed genes (DEGs) for the 17 mM glucose spike experimental condition with |log2 FC|≥ 1.5 and adjusted *P* value ≤ 0.05 as the threshold. The red dots represent 43 significantly upregulated genes, and the blue dots represent 42 significantly downregulated genes. **(B)** Venn diagram illustrating the number of unique and shared DEGs from the 10 mM, 17 mM, and 50 mM glucose challenge experimental conditions. **(C)** Summary of the significantly downregulated genes for the 17 mM glucose spike experimental condition assigned to functional classes according to GO and KEGG pathways.

### *S. epidermidis* Transcription Response to a 20-minute 17 mM Glucose Spike

We identified 85 of 2354 genes (∼4% of the genome) that change in response to a 17 mM glucose spike with log2FC values ≥ 1.5, among which 43 were upregulated and 42 were downregulated (Table S6). Downregulated and upregulated genes were expressed over a −5 to +5 log2FC range (Figure 4A). While gene expression changes are similar across all glucose levels, we observed a more robust change (i.e., −5 to +5 log2FC), a higher number of upregulated genes, and a higher total number of DEGs in the 17 mM glucose condition (Table S3-S5; Table S7).

Among the most downregulated genes (**∼**6-34-fold) in the 17 mM glucose spike condition were *pflB* and members of the *glpR-pfkB* operon, which plays an essential role in the utilization of fructose, (F1613_RS05150 (DeoR/GlpR family DNA-binding transcription regulator), *pfkB*, and F1613_RS05160 (PTS fructose transporter subunit IIABC)) (Ge et al., 2024) (Table S6). We found that L-serine ammonia-lyase iron-sulfur-dependent subunits alpha and beta (*sdaAA* and *sdaAB*), *raiA*, F1613_RS01060 (PTS sugar transporter subunit IIC), and F1613_RS00520 (nitrate reductase subunit alpha) were also downregulated (**∼**6-10-fold) (Table S6). Six hypothetical proteins and one uncharacterized gene (seven total) were downregulated in the 17 mM glucose spike condition. tRNA genes accounted for almost 60% (24 of 43) of the upregulated genes in the 17 mM glucose spike condition, consistent with increased protein synthesis and faster growth rates in the presence of glucose (Halsey et al., 2017). F1613_RS07200 (solute carrier family 23 protein) and *ffs* were among the most upregulated genes (**∼**7 to 11-fold) in the 17 mM glucose spike condition. Two hypothetical proteins and three uncharacterized genes (five total) were also upregulated.

### Functional Classification of Downregulated Genes in *S. epidermidis* in Response to a 17 mM Glucose Spike

To further understand the functions of significantly downregulated genes we used the data from the 17 mM glucose spike condition to assign functional pathways against the GO and KEGG databases. We ordered pathways based on increasing significance level (*P* value) (Figure 4C). Functional pathways with decreased expression include Carbohydrate Metabolism, Butanoate Metabolism, TCA Cycle, Propanoate Metabolism, Lipoic Acid Metabolism, Carbohydrate Transport, Hexose Metabolism, Oxidative Phosphorylation, Phosphoenolpyruvate-Dependent Sugar Phosphotransferase system (PTS), and Amino Acid Metabolism (Figure 4C; Table S6). We observed several downregulated pathways likely consistent with carbon catabolite repression (CCR) (Görke and Stülke, 2008).

### *S. epidermidis* Transcription Response to Persistent Glucose via Batch Culture

To identify differentially expressed genes in *S. epidermidis* exposed to persistent glucose via batch culture, we grew cells overnight in medium containing 0.2% w/v or 1% w/v glucose. We used RNA sequencing to analyze gene expression profiles and compared the expression profiles of cells exposed to 1% w/v glucose against cells exposed to 0.2% w/v glucose. Differentially expressed genes (DEGs) of significance were defined using the following criteria: |log2 fold change| (i.e., log2FC) ≥ 1.5 and adjusted *P* value < 0.05. By these criteria, we identified 195 of 2354 genes (∼8% of the genome) with log2FC values ≥ 1.5, among which 133 were upregulated and 62 were downregulated (Table S8). We observed more upregulated genes, a higher total number DEGs, and unique gene expression changes in the persistent glucose via batch culture experimental condition compared to the 20-minute glucose spike experimental condition (Table S3-S7; Table S8).

Among the most upregulated genes (∼13-30-fold) in the persistent glucose condition were members of the *nrdDG* operon (*nrdD* and *nrdG*), which encodes for an oxygen-independent ribonucleotide reductase (Masalha et al., 2001), and the *dha* operon (F1613_RS03960 (glycerol dehydrogenase), *dhaK*, *dhaL*, *dhaM*)), which encodes for components of the glycerol dehydrogenase- and PTS-dependent dihydroxyacetone kinase system (Céline Monniot et al., 2012) (Table S8). Genes involved in nitrate/nitrite reduction (*narGHJI*, *nirBD*, *nreABC, and* F1613_RS00485 (NarK/NasA family nitrate transporter)) were also upregulated (**∼**4.8-11.9-fold) (Kamps et al., 2004) (Table S8). Sixteen genes involved in glycolysis, gluconeogenesis, and the TCA cycle including the glycolytic *gapA* operon (*gap*, F1613_RS05590 (phosphoglycerate kinase), *tpiA*, *gpmI*, and *eno*), the *alsS/budA* operon, F1613_RS00620 (2,3-diphosphoglycerate-dependent phosphoglycerate mutase), F1613_RS01410 (fructose bisphosphate aldolase), *fdaB*, F1613_RS01355 (L-lactate dehydrogenase), *sdaAA*, *pyk*, *ilvB*, F1613_RS06110 (glucose-6-phosphate isomerase) and *sdhB* were slightly upregulated (∼3-8 fold) in the persistent glucose condition, consistent with previous observations of glucose-responsive genes in *S. aureus* (Seidl et al., 2009). Seven hypothetical proteins were also upregulated (Table S8).

We observed downregulation (up to ∼7 fold) of the energy-coupling factor (ECF) transporter module components (F1613_RS11970 (energy-coupling factor transporter ATPase), F1613_RS11965 (energy-coupling factor transporter ATPase), F1613_RS11960 (energy-coupling factor transporter transmembrane protein EcfT)) (Slotboom, 2013), F1613_RS03610 (isoprenylcysteine carboxyl methyltransferase family protein), and *ugpC* (Table S8). F1613_RS05940, *dltC*, and *dltD*, components of the *dlt* operon required for the d-alanylation of teichoic acids in gram-positive bacterial cell walls (Kovacs et al., 2006), were also downregulated (∼3-4 fold). We observed downregulation of four transcription regulators including *rsp*, F1613_RS01465 (GbsR/MarR family transcription regulator), F1613_RS08735 (AraC family transcription regulator), and F1613_RS10440 (helix-turn-helix transcription regulator) by 3.3-, 3.5-, 3.7-, and 4.2-fold, respectively (Table S8). Two hypothetical proteins were also downregulated in the persistent glucose condition (Table S8).

### *S. epidermidis* Transcription Response to a Step Down in Glucose Concentration from 10 mM to 2 mM

To identify differentially expressed genes in *S. epidermidis* exposed to a step down in glucose concentration, we challenged mid-exponential phase cells by subjecting them to a 10 mM glucose spike for 20 minutes immediately followed by a 2 mM glucose spike for 20 minutes (Figure 1B). We used RNA sequencing to analyze gene expression profiles and compared the expression profiles of cells exposed to a step down in glucose concentration against cells exposed to a 10 mM glucose spike only. Differentially expressed genes (DEGs) of significance were defined using the following criteria: |log2 fold change| (i.e., log2FC) ≥ 1.5 and adjusted *P* value ≤ 0.05. By these criteria, we identified 43 of 2354 genes (∼1.8% of the genome) with log2FC values ≥ 1.5, among which 10 were upregulated and 33 were downregulated (Table S9; Figure S5). Downregulated and upregulated genes were expressed over a −6 to +3 log2FC range (Figure 5A).

**Figure 5.**
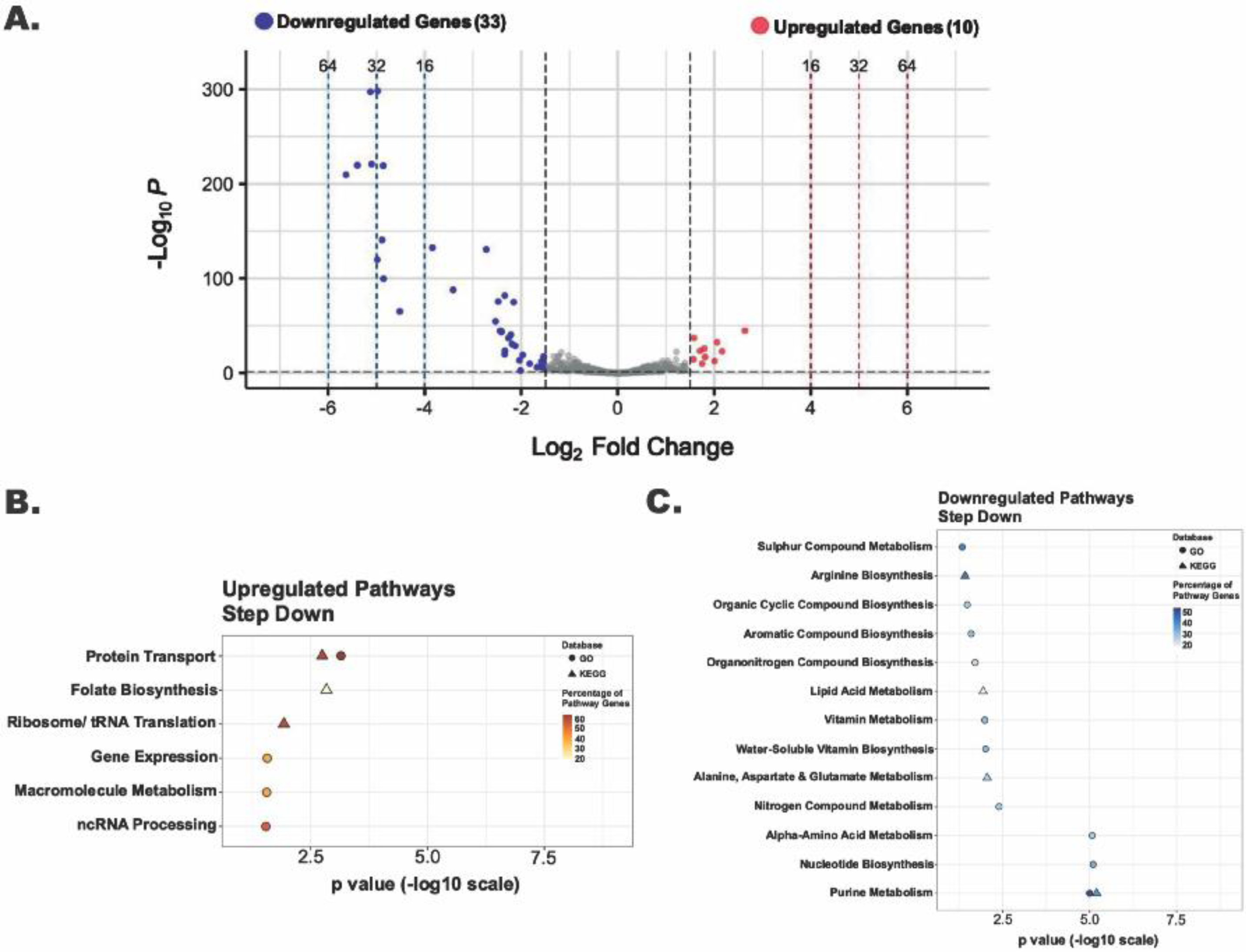
Genes Involved in Purine Metabolism Are Significantly Downregulated in Response to a Step Down in Glucose Concentration From 10 mM to 2 mM. **(A)** Volcano plot showing the differentially expressed genes (DEGs) for the Step-down experimental condition with |log2 FC|≥ 1.5 and adjusted *P* value ≤ 0.05 as the threshold. The red dots represent 10 significantly upregulated genes, and the blue dots represent 33 significantly downregulated genes. Summary of the significantly upregulated **(B)** and downregulated **(C)** genes for the Step-down experimental condition assigned to functional classes according to GO and KEGG pathways.

We observed upregulation (**∼**3-6-fold) of F1613_RS03760 ((NAD(P)-binding domain-containing protein), *betB*, *betA*, F1613_RS03755 (nucleoside recognition domain-containing protein), *rpsN*, F1613_RS06020 (NAD(P)-binding domain-containing protein), F1613_RS00615 (putative metal homeostasis protein), F1613_RS02245 (putative sulfate exporter family transporter), F1613_RS03765 (zinc ABC transporter substrate-binding protein), and F1613_RS01245 (aminotransferase class I/II-fold pyridoxal phosphate-dependent enzyme) in the step-down experimental condition. Among the most downregulated genes (∼5-50-fold) were members of the purine biosynthetic operon (*purEKCSQLFMNHD*), which encodes for 11 enzymes that convert phosphoribosyl pyrophosphate (PRPP) to inosine-5′-monophosphate (IMP) (Goncheva et al., 2019), purine biosynthesis-associated gene *purB*, and glycine cleavage system genes (*gcvT*, *gcvPA*, *gcvPB*). One hypothetical protein was also downregulated in the step-down experimental condition (Table S9).

### Functional Classification of Differential Expressed Genes in *S. epidermidis* in Response to a Step Down in Glucose Concentration from 10 to 2 mM

We used the data from the step-down experimental condition to assign functional pathways against the GO and KEGG databases. We ordered pathways based on increasing significance levels (*P* value) (Figure 5C). Functional pathways with decreased expression include Purine Metabolism, Nucleotide Biosynthesis, Amino Acid Metabolism, Nitrogen Compound Metabolism, Vitamin Metabolism, Lipid Acid Metabolism, Organic Compound Biosynthesis, and Sulphur Compound Metabolism (Figure 5C; Table S9). Among upregulated pathways Protein Transport scored the highest significance, according to both GO and KEGG pathway enrichment analysis, under the step-down experimental condition (Figure 5B).

We constructed a Venn diagram to understand the relationship between our step-down, 10 mM glucose spike (G10), and 2 mM glucose spike (G2) data sets (Figure S5); we observed no shared differentially expressed genes (DEGs) in common among the step-down condition (from 10 to 2 mM glucose) and G10 (from 0 to 10 mM glucose). There were also no shared differentially expressed genes among the step-down (from 10 to 2 mM glucose) and G2 (from 0 to 2 mM glucose) experimental conditions either, indicating potentially unique gene expression changes as a function of increasing versus decreasing glucose concentrations (Figure S5; Table S3; Table S9).

We sought to further understand if and how genes might be differentially expressed at an intermediate glucose concentration (2 mM glucose) as a function of whether cells had been previously exposed to a lower (0 mM) or higher (10 mM) glucose concentration. If prior glucose concentrations do not matter, we would expect no such differences. We performed scatter plot analysis of expression levels for all genes at 2 mM glucose as a function of prior glucose concentration (Figure 6). Most genes differentially expressed under a 0 to 2 mM glucose spike were similarly expressed under a 10 to 2 mM glucose step down (Figure 6 blue dots). Over 14 genes differentially expressed under a 10 to 2 mM glucose step down were not similarly expressed under a 0 to 2 mM glucose spike (Figure 6 red dots; Discussion). Further analysis indicated these genes are primarily involved in purine metabolism (above; Table S9).

**Figure 6.**
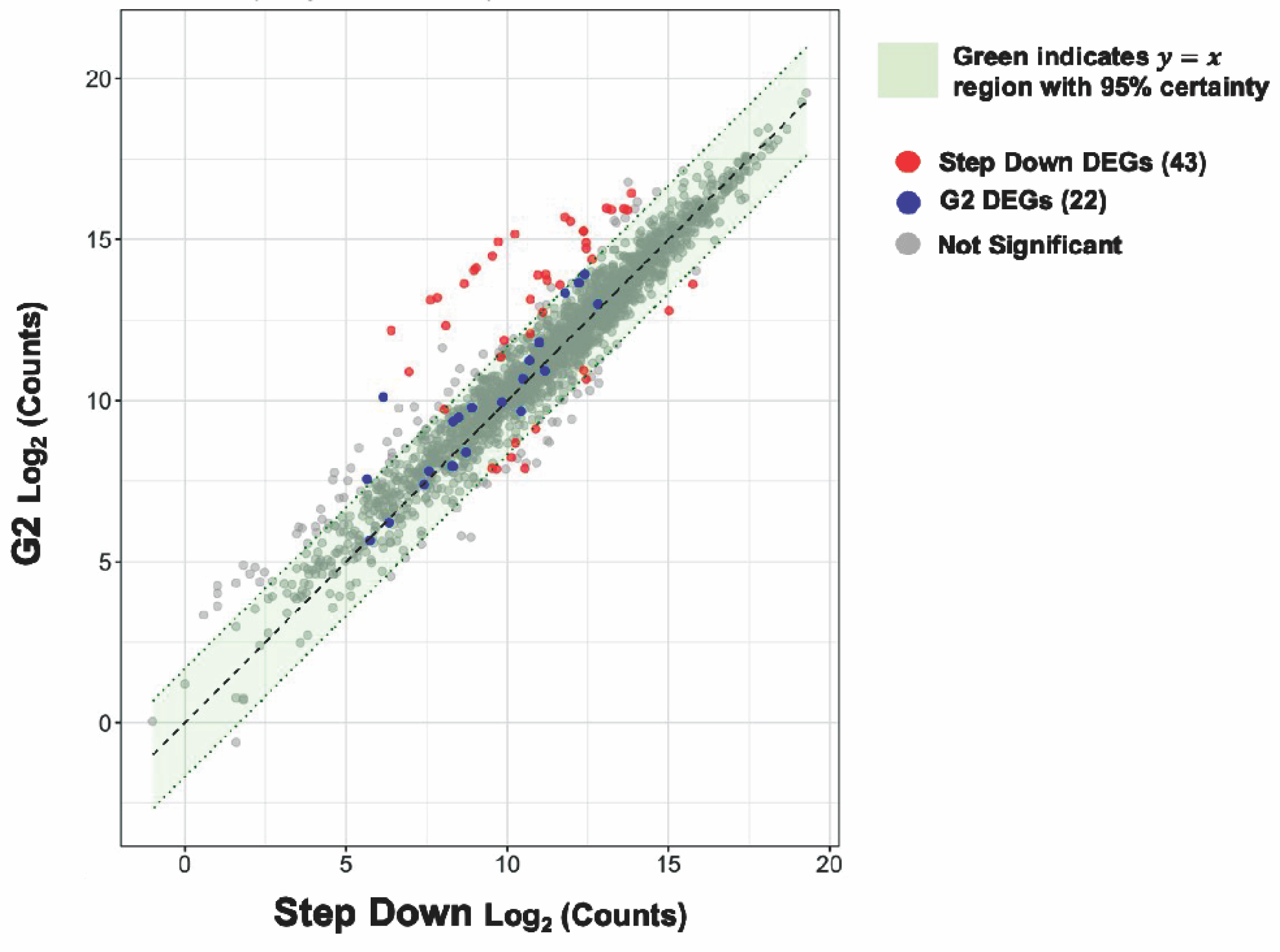
Expression levels of purine biosynthesis genes at intermediate glucose levels are sensitive to prior glucose levels. Scatter plot visualizing the relationship between the step-down and 2 mM glucose spike (G2) experimental conditions. Each dot denotes a single gene. The red and blue dots represent step-down, and G2 differentially expressed genes (DEGs) respectively. The gray dots represent genes with no significant change. A 95% confidence interval was calculated around the residuals of gene expression differences between the two experimental groups. Genes that fall within the green highlighted region are predicted to have near identical average expression levels with 95% certainty.

### Discriminating Between Glucose and Heat Shock Conditions

Differential gene expression analysis of and within the skin microbiome might be useful as a potential platform for clinical diagnosis. To explore this idea, we compared gene expression levels during heat shock to those observed during high (17 mM) glucose levels. Most (∼93.6%) genes are similarly expressed (95% c.i.) under both conditions (Figure 7). However, 341 and 60 genes are differentially expressed under heat shock or high glucose, but not both conditions, respectively. Such genes may offer a starting point for developing nucleic acid amplification-based methods for determining the current or prior physical experience of microbes on patients.

**Figure 7.**
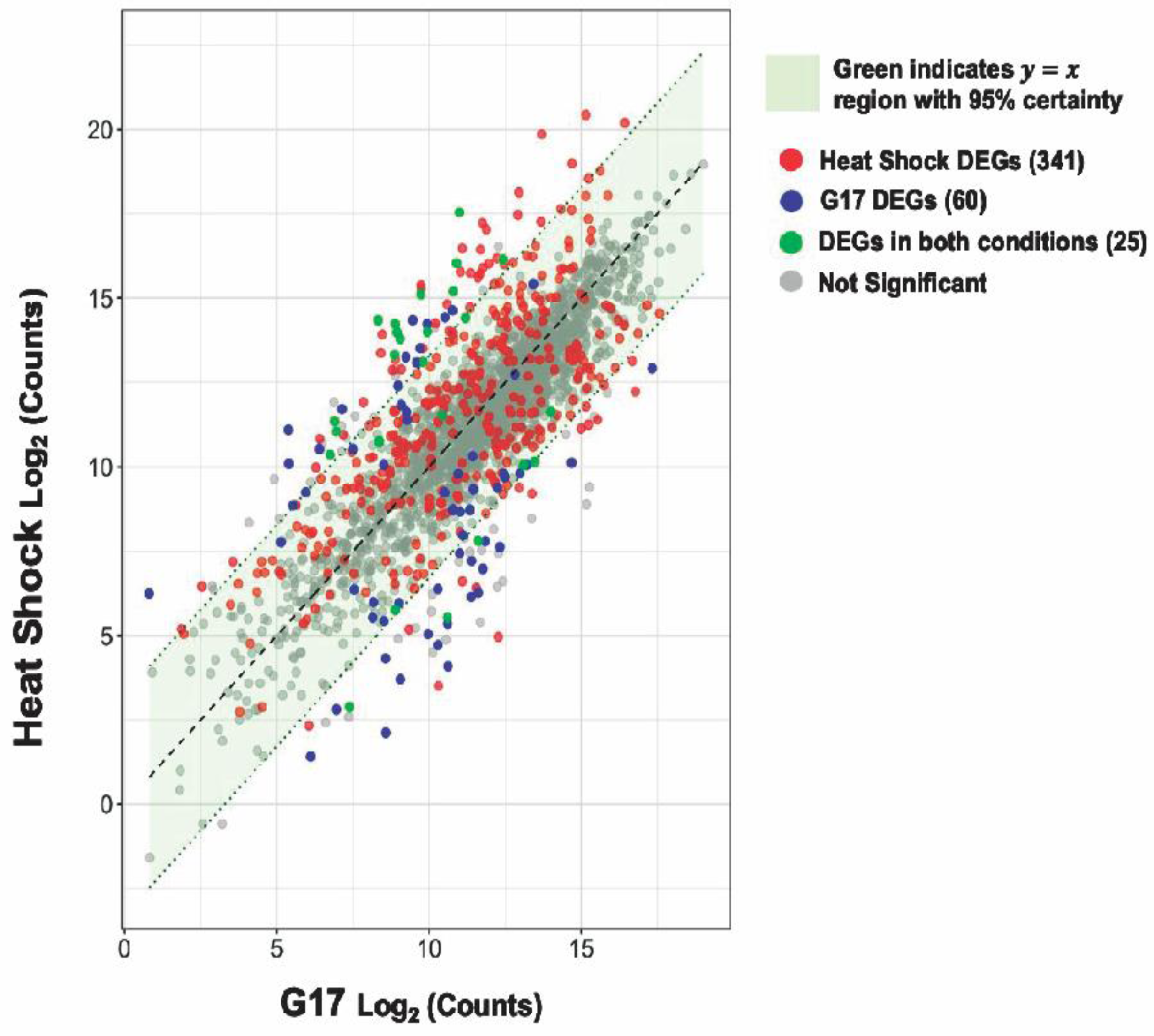
Heat shock and glucose spikes create statistically unique signatures. Scatter plot visualizing the relationship between the heat-shock and 17 mM glucose spike (G17) experimental conditions. Each dot denotes a single gene. The red and blue dots represent heat-shock, and G17 differentially expressed genes (DEGs) respectively. The green dots represent DEGs found in both conditions and the gray dots represent genes with no significant change. A 95% confidence interval was calculated around the residuals of gene expression differences between the two experimental groups. Genes that fall within the green highlighted region are predicted to have near identical average expression levels with 95% certainty.

## DISCUSSION

To support bioengineering of skin microbes to diagnose, monitor, or treat disease, we sought to understand how *S. epidermidis* responds to environmental perturbations including heat shock and medically relevant glucose levels. We used RNA sequencing to investigate differential gene expression followed by gene set enrichment analysis (GSEA) to understand the functions of differentially expressed genes. We observed an *S. epidermidis* heat-shock response that mostly resembles the heat-shock response of related staphylococcal species and other gram-positive bacteria (below). We observed carbon catabolite repression in response to a range of glucose spikes, upregulation of genes involved in glycolysis, gluconeogenesis, and the TCA cycle in response to persistent glucose via batch culture, as well as a potentially unique gene expression signature in response to a step down in glucose concentration from 10 to 2 mM. Building upon our analyses we curated a subset of glucose-responsive genes that might serve as starting points for engineering endogenous dynamic control of circuits in *S. epidermidis*.

We observed contrasting patterns of gene expression depending on whether cells were exposed to a spike or persistent level of glucose. For example, we observed downregulation (up to ∼34 fold) across all five glucose spike conditions for genes involved in lactose metabolism, ribose transport, fructose utilization, proline catabolism, the glyoxalase pathway, the succinate dehydrogenase complex, and ethanol degradation (Table S3-S7). We believe this repression of genes involved in secondary carbon source utilization to be convincing evidence of carbon catabolite repression (CCR) in our glucose spike data (Görke and Stülke, 2008). By contrast, we found no evidence of CCR in our persistent glucose via batch culture data. (Table S8). As a second example, while we observed the induction (∼3-8 fold) of several essential glycolytic genes, the *dha* operon, gluconeogenesis genes, and TCA cycle genes in our persistent glucose via batch culture samples (Table S8), we did not observe such gene expression patterns among the upregulated genes in our glucose spike data. Instead, tRNA genes accounted for most of the upregulated genes in our glucose spike data (Table S3-S8). One explanation could be that *S. epidermidis* first adapts to glucose exposure by preferentially downregulating genes involved in secondary carbon source utilization to avoid the production of proteins that are not useful in the presence of glucose; only following sufficiently prolonged exposure to glucose does *S. epidermidis* adjust its transcriptome to upregulate genes involved in glucose utilization. We note that Seidl et al. found in *S. aureus* that a 30-minute exposure to 10 mM glucose was sufficient to realize gene expression changes similar to our prolonged exposure conditions, suggesting that between 20 to 30 minutes could be sufficient to fully transition to a persistent glucose transcriptome in *S. epidermidis* (Seidl et al., 2009).

Under heat shock conditions we found patterns of gene expression similar to other *Staphylococcus* species. For example, at 45°C, we observed upregulation of F1613_RS04215 (*CtsR* family transcription regulator) and *hrcA* (Table S1), known heat-shock gene expression regulators in *Staphylococcus aureus, Bacillus subtilis*, and other firmicutes (Derre, Rapoport, and Msadek, 1999; Chastanet, Fert, and Msadek, 2003; Schumann, 2003). We also observed rapid induction of *clpB*, *clpP*, and the *dnaK*, *groESL*, and *clpC* operons (Table S1). Our data also provides evidence of an *S. epidermidis* heat-shock regulatory network that utilizes both the *hrcA-* and *ctsR-*encoded repressors. For example, we carried out DNA motif analysis and found CtsR (GGTCAAA/T) and CIRCE (controlling inverted repeat of chaperone expression) operator sequences arranged in tandem upstream of the *dnaK* and *groESL* operons consistent with previous observations of dual heat-shock regulation by HrcA and CtsR in *S. aureus* and *S. epidermidis* (Derre, Rapoport and Msadek, 1999; Chastanet, Fert, and Msadek, 2003) (Figure S6). We also found CtsR recognition sequences upstream of *clpB*, *clpP*, and the *clpC* operon also consistent with previous observations of CtsR regulons in *B. subtilis* and *Streptococcus pneumoniae* (Derre et al., 1999; Chastanet et al., 2001) (Figure S6).

While we observed upregulation of universal stress proteins (F1613_RS09680 and F1613_RS09700), we did not detect upregulation of the general stress-responsive alternative sigma factor *sigB,* which is a component of the heat-shock regulon in *S. aureus, B. subtilis*, and *Listeria monocytogenes* (Kullik and Giachino, 1997; Schumann, 2003; Ferreira, O’Byrne, and Boor, 2001). By contrast, we did observe upregulation (∼5 fold) of F1613_RS09995, another sigma-70 family RNA polymerase sigma factor (Table S1). This difference suggests that the *S. epidermidis* heat-shock regulatory network may differ slightly from that of *S. aureus* and other gram-positive bacteria.

We compared the genome-wide *S. epidermidis* heat-shock response to the 17 mM glucose spike (G17) and step-down responses (Figure 8). We observed a more robust increase in gene expression in response to heat shock (i.e., −4 to +6 log2FC range) compared to G17 (i.e., −5 to +5 log2FC) and step down (i.e., −6 to +3 log2FC range) and detected more differentially expressed genes (DEGs) in the heat-shock condition (366 genes) compared to G17 (85 genes) and step-down conditions (43 genes) (Figure 2A; Figure 4A; Figure 5A). In response to acute heat stress and subsequent loss of protein homeostasis (e.g., due to heat-induced protein denaturation, misfolding, and aggregation), we observed a rapid and global reprogramming of gene expression, unlike the transcription changes observed when *S. epidermidis* adapts to a preferred carbon source (e.g., glucose) at non-toxic concentrations (Figure 8; Figure 3A). We believe these disparate gene expression profiles could be of limited clinical utility; more specifically, DEGs unique to heat shock (341 genes) or high glucose (60 genes) may be a promising starting point for the development of simple nucleic acid-based tools for the diagnosis and monitoring of disease (Figure 7).

**Figure 8.**
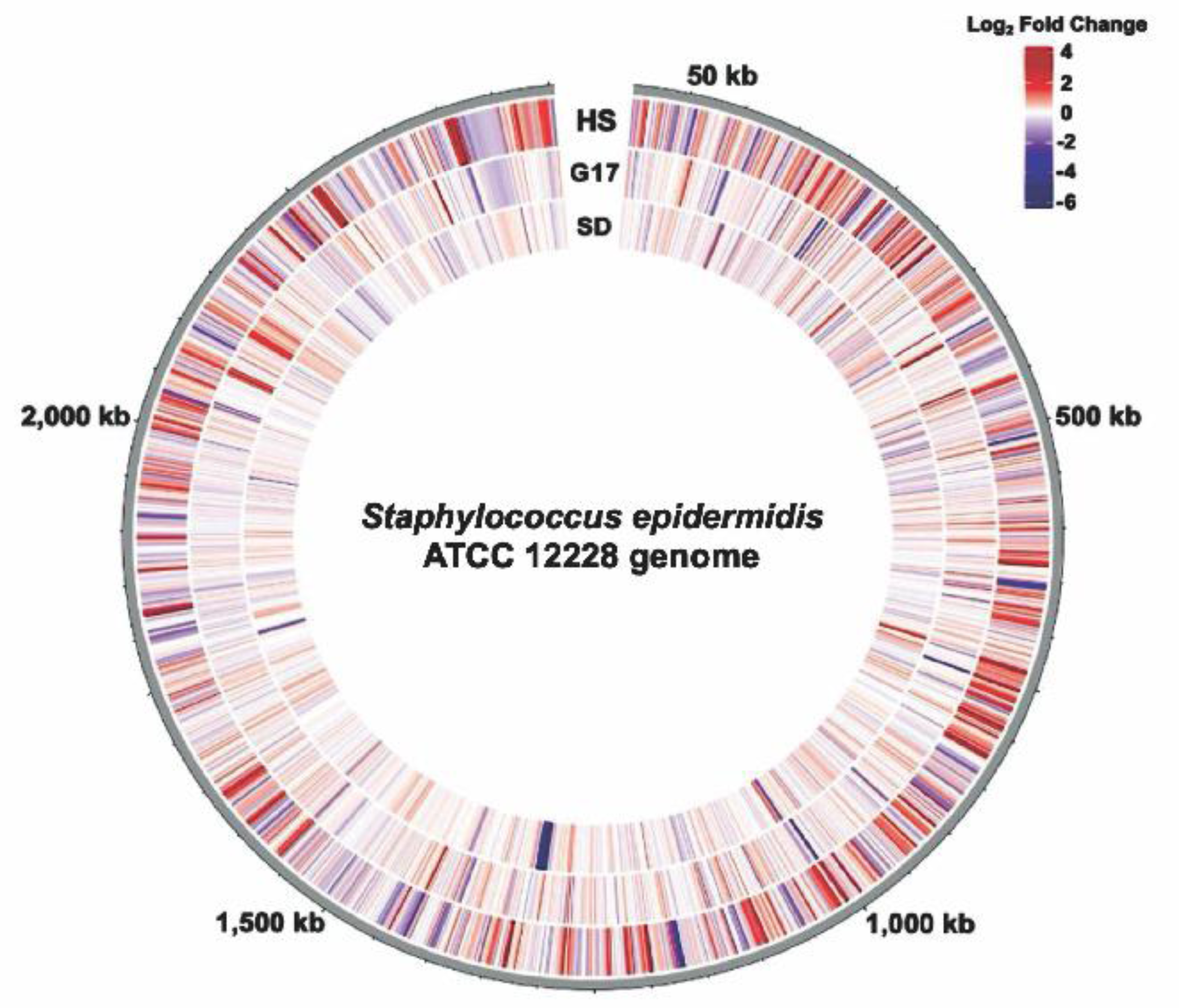
The Genome-wide Transcription Response of *Staphylococcus epidermidis* to Perturbations. Circular transcriptome map showing normalized gene expression levels in the *Staphylococcus epidermidis* genome. Log2-fold change relative to control for cells exposed to Heat Shock (HS), a 17 mM glucose spike (G17), or Step Down (SD) experimental conditions. Each bar denotes a single gene. The red bars represent significantly upregulated genes, and the blue bars represent significantly downregulated genes.

We observed downregulation (up to ∼50-fold) of genes involved in purine biosynthesis (*purEKCSQLFMNHD*) in response to a step down in glucose concentration from 10 to 2 mM (Table S9). We did not observe such downregulation in the G2 (from 0 to 2 mM glucose) or G10 (from 0 to 10 mM glucose) glucose spike conditions (Table S3; Table S5). Further, we found no differentially expressed genes (DEGs) in common among the step-down and G10 conditions and the step-down and G2 conditions (Figure S5). Taken together we wondered if there is a unique step-down gene expression signature that does not resemble that of G2 or G10. We performed scatter plot analysis to visualize the relationship between the step-down and G2 conditions (Figure 6). We noticed that, while most genes are similarly expressed under both conditions, over 14 genes differentially expressed under step-down conditions were not similarly expressed under G2 conditions (Figure 6, red dots). Further analysis revealed that these genes were mainly involved in purine biosynthesis. We note that our step-down samples underwent two rounds of centrifugation while our G10 samples underwent a single round of centrifugation prior to RNA harvesting (Methods); this methodological difference may account for the unique step-down gene expression signature observed here.

Finally, we sought to identify glucose-responsive promoters that might eventually be used to control the expression of an insulin gene in a bioengineered *S. epidermidis* strain developed to aid in treating diabetes. To this end, we constructed glucose concentration-response curves across medically relevant (G2-G17) glucose levels. We identified 38 glucose-responsive genes that might serve as ON or OFF switches for controlling synthetic genetic systems (Figure S3; Figure 3B). Most (∼70%) of the potential switches that exhibited an OFF-to-ON transition were tRNA genes (Figure S3). We suspect these switches are not specific to glucose given that increased tRNA expression might also occur in response to various other carbon sources (Dong, Nilsson and Kurland, 1996). We also observed 19 potential ON-to-OFF switches (Figure S3). Each glucose-responsive gene reported here is a starting point requiring additional characterization (e.g., response specificity) to identify those most appropriate for any given application (e.g., controlling expression of insulin in a glucose-dependent manner).

The human skin microbiome is a diverse and dynamic microbial community that plays an essential role in maintaining our health and well-being. A more intimate understanding of how our skin microbes adapt to environmental perturbations (e.g., stress or increased glucose levels) is required to ultimately enable development of bioengineered skin microbes that can help diagnose and treat disease. We hope our investigation of the genome-wide transcription response in *S. epidermidis* to heat shock and medically relevant glucose concentrations helps further motivate ongoing work. We are excited to imagine a future in which the bioengineering of skin microbes has been made routine, helping doctors and patients to realize healthier lives and better clinical outcomes.

## AUTHOR CONTRIBUTIONS

KNB: conceptualization, design and execution of the experiments, data interpretation, funding acquisition and writing (original draft, review, and editing). AG: data analysis and interpretation, data visualization, writing (part of the methods section, and review). RN: data preprocessing pipeline development and writing (part of the methods section, and review). DE: supervision, data interpretation, funding acquisition, and writing (review, and editing). All authors contributed to the article and approved the final manuscript.

## ACKNOWLEDGEMENTS

We thank John Glass, Richard Gallo, and Yo Suzuki for their support, mentorship, and helpful comments.

---

**Figure S1.**
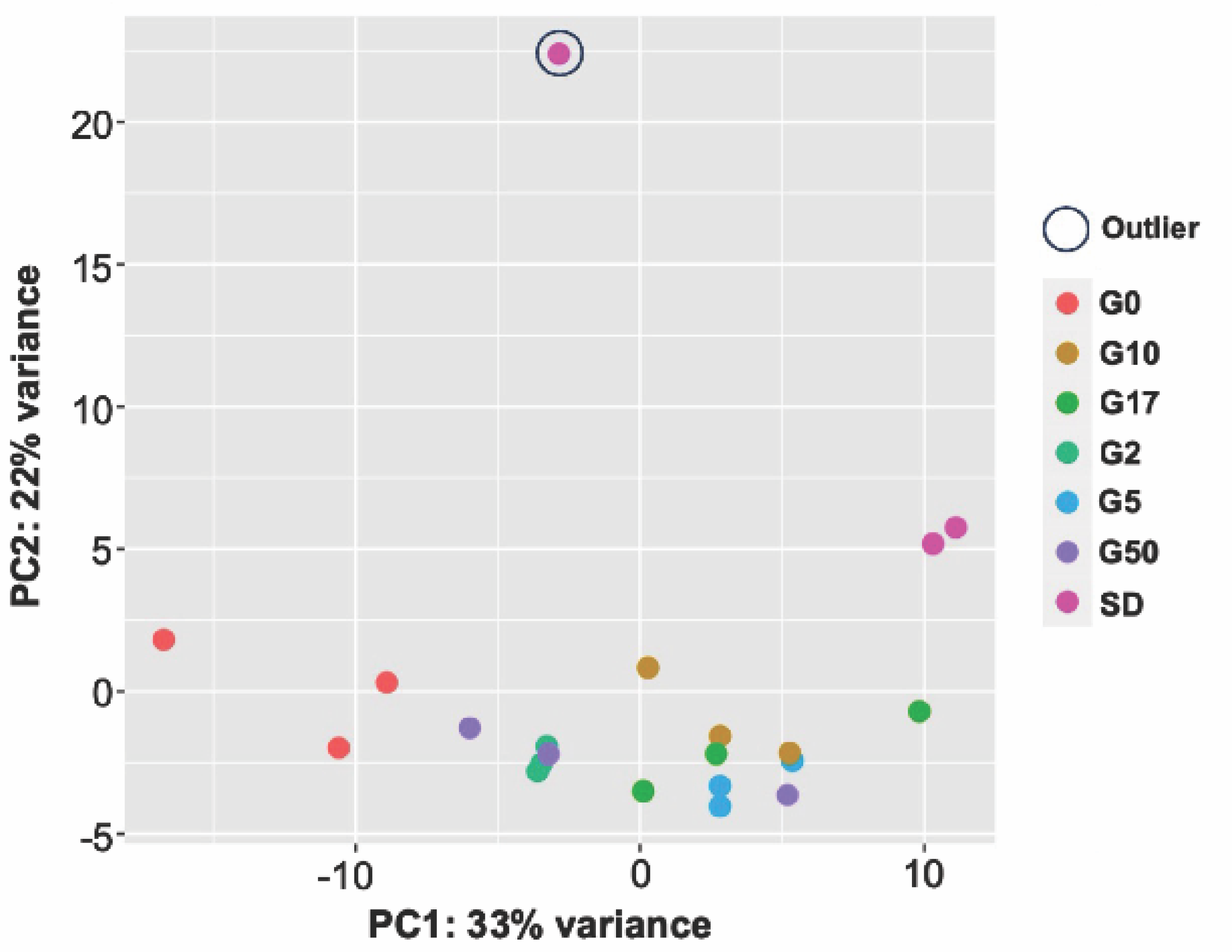
Principal Component Analysis (PCA) Reveals One Step-Down Replicate is an Outlier. PCA plot of RNA-seq data for three biological replicates for the Glucose Challenge and Step-Down experimental conditions.

**Figure S2.**
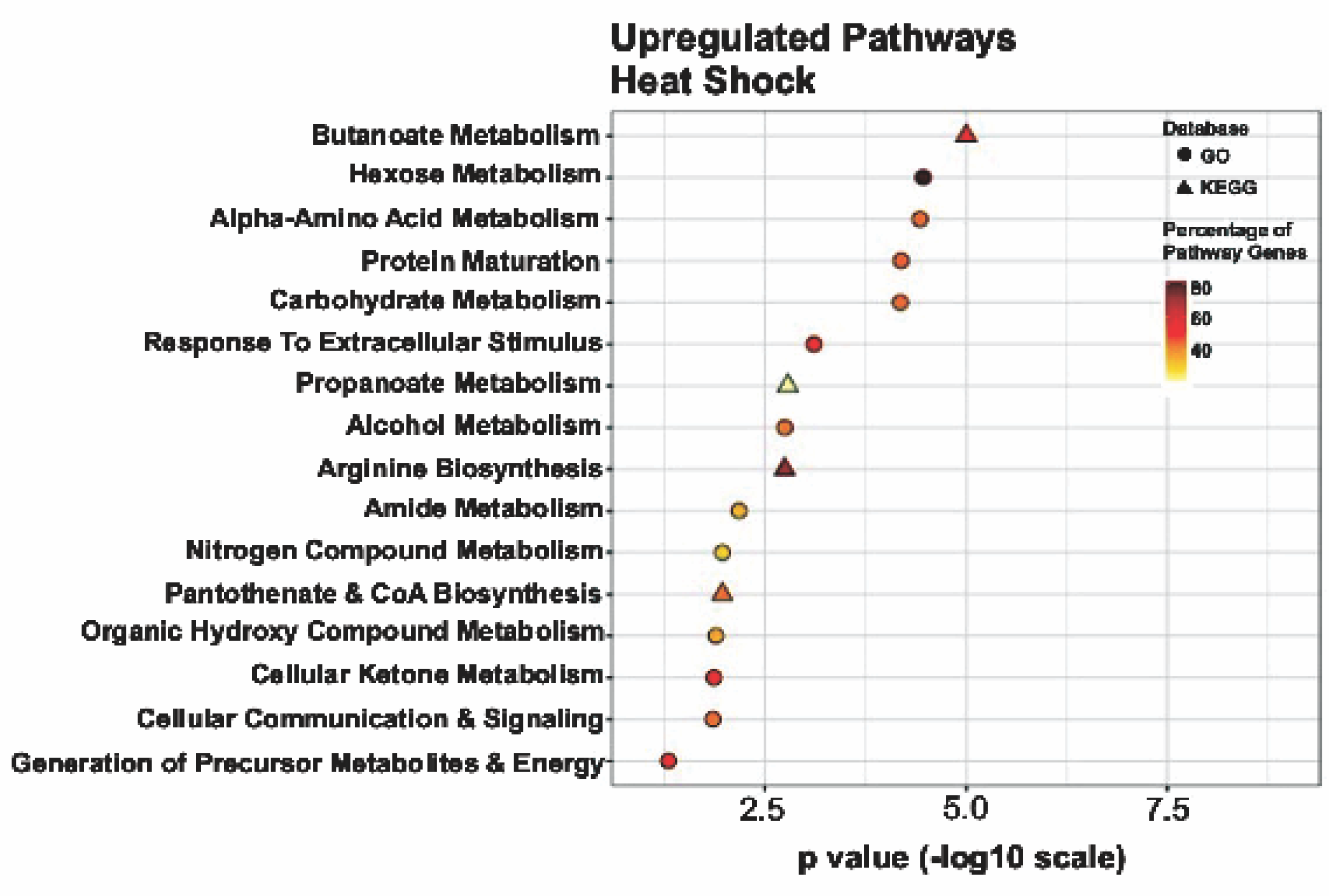
Genes Upregulated During Heat-shock Span Metabolism and Cell Signaling. Summary of the significantly upregulated genes for the heat-shock experimental condition assigned to functional classes according to GO and KEGG pathways.

**Figure S3.**
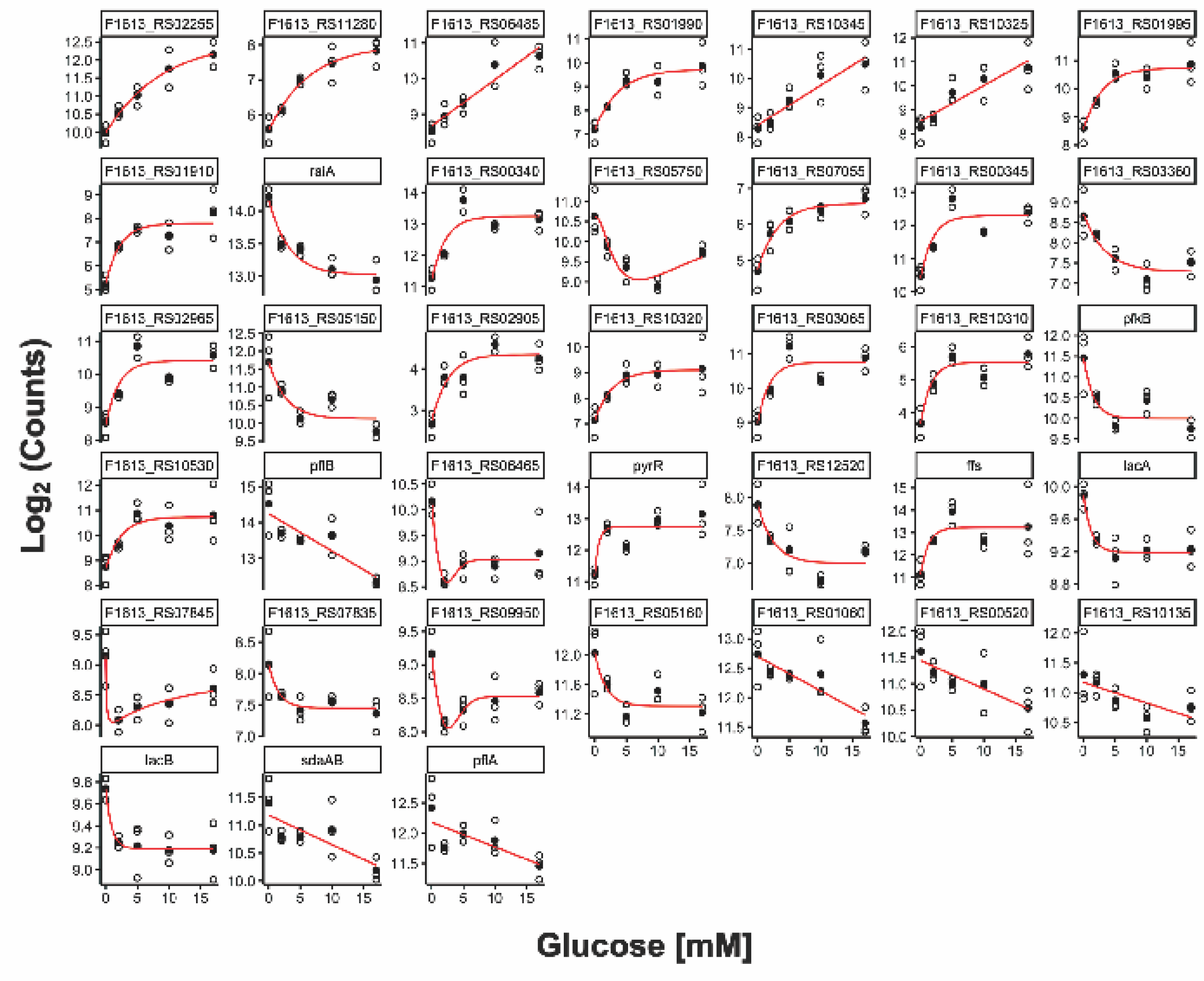
Thirty-eight Candidate Genes that Have Potentially Interesting Glucose-Responsive ON or OFF Switch Properties. Concentration-response curves generated for genes with absolute log2 fold change values ≥ 2 in at least one medically relevant (G2-G17) glucose challenge experimental condition.

**Figure S4.**
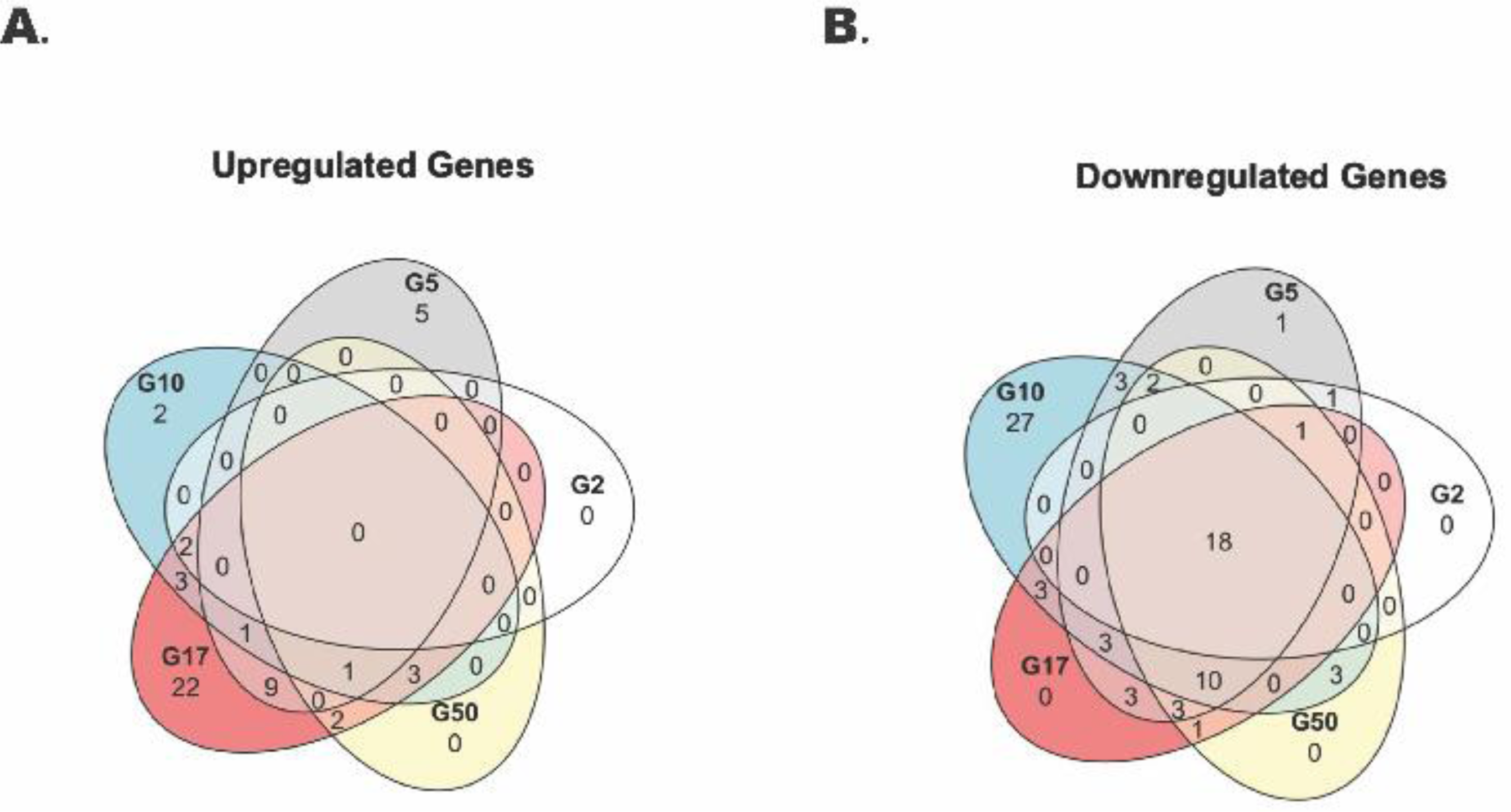
Twenty-eight Genes are Downregulated in Common in Response to Increasing Glucose Spike Concentrations. Venn diagrams illustrating the number of unique and shared upregulated **(A)** and downregulated genes **(B)** from the 2 mM (G2), 5 mM (G5), 10 mM (G10), 17 mM (G17), and 50 mM (G50) glucose challenge experimental conditions.

**Figure S5.**
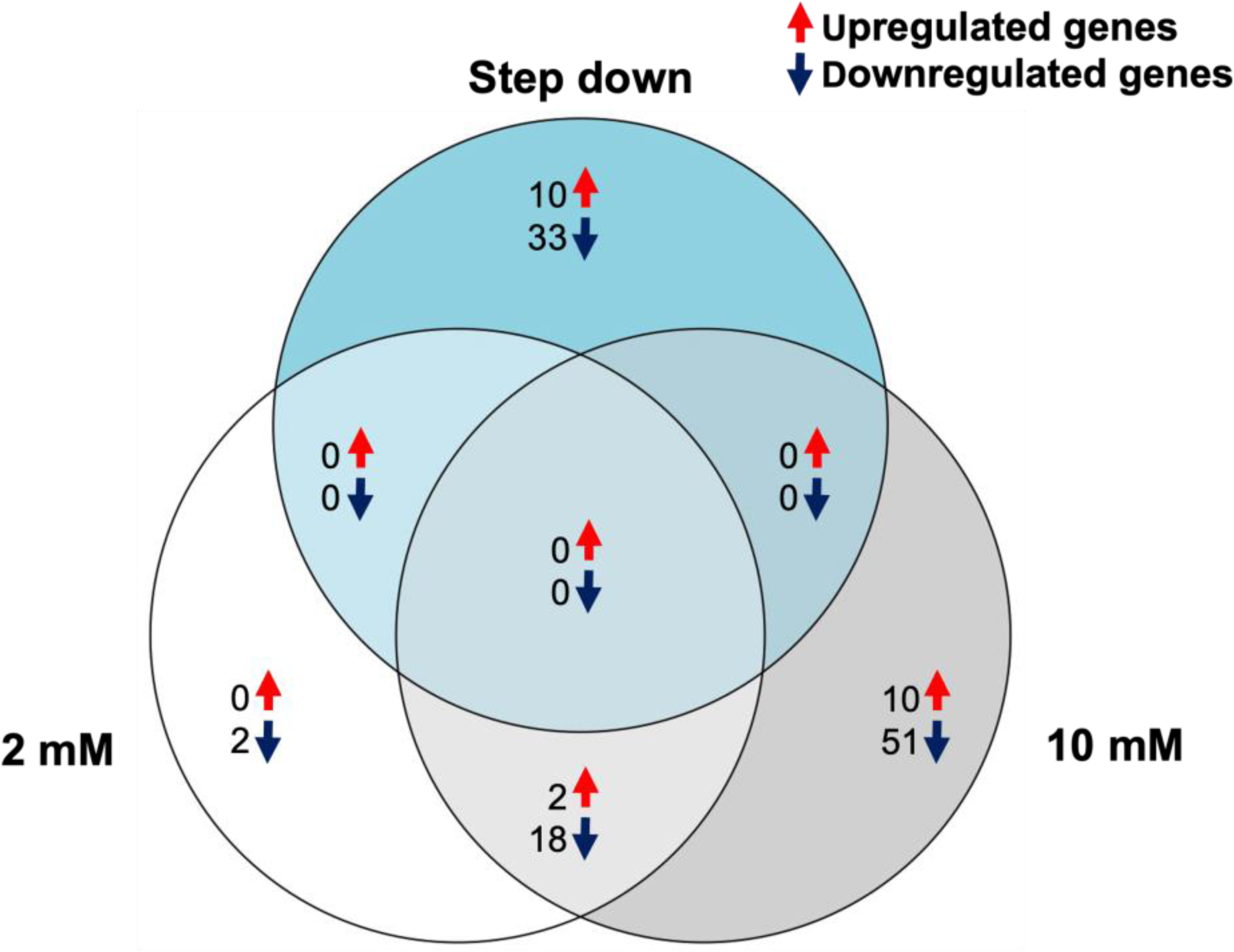
Step-down Samples Have a Unique Gene Expression Profile Distinct from the 2 mM and 10 mM Glucose Spike Profiles. Venn diagram illustrating the number of unique and shared DEGs from the 2 mM, 10 mM, and step-down experimental conditions.

**Figure S6.**
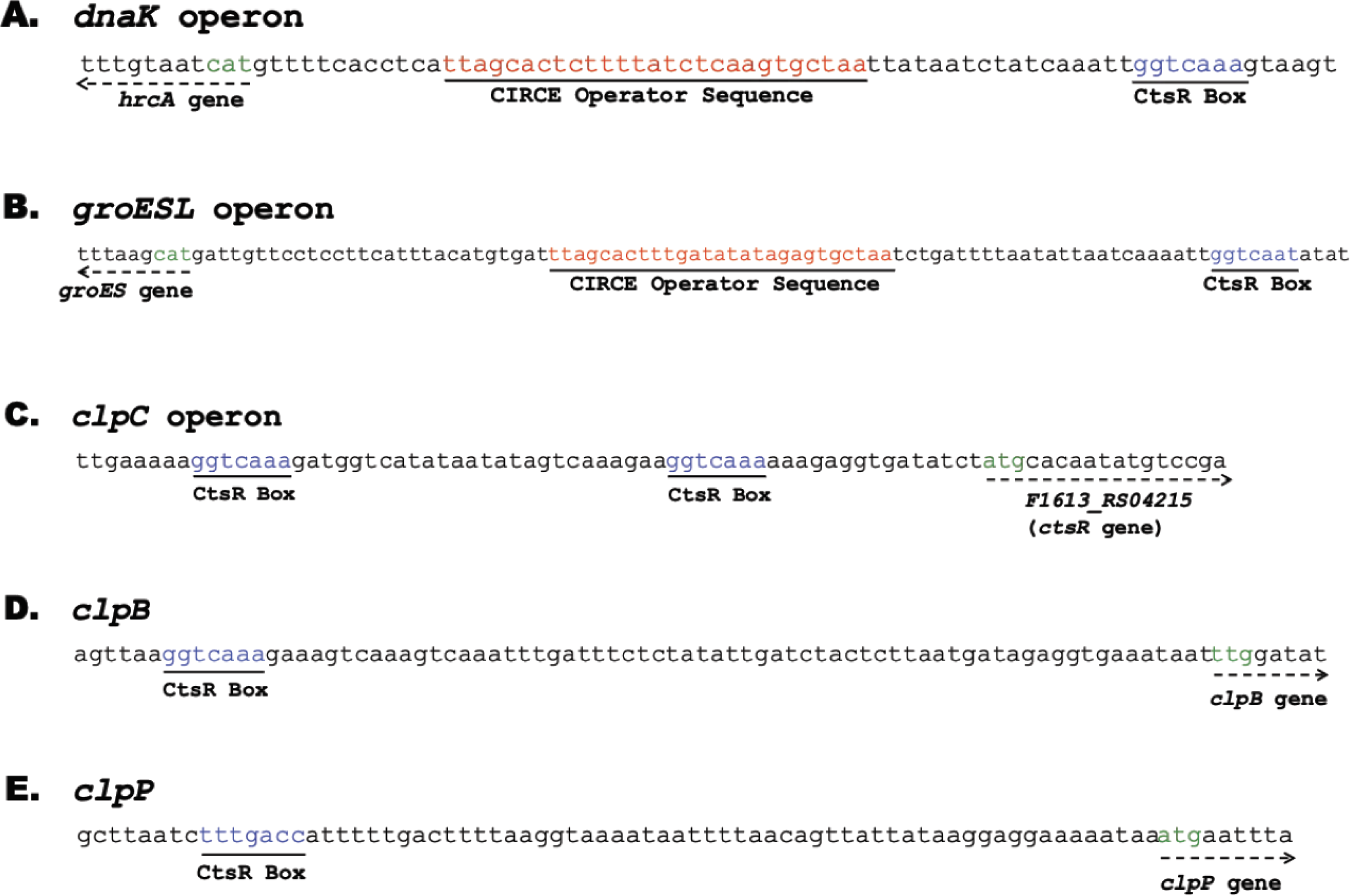
DNA Motif Analysis Revealed CtsR and CIRCE Operator Sequences Arranged in Tandem Upstream of dnaK and groESL Operons. CIRCE (TTAGCACT-N11-AGTGCTAA) (red) and/or CtsR (GGTCAAA/T) (blue) operator sequences arranged upstream of the **(A)** *dnaK* operon, **(B)** *groESL* operon, **(C)** *clpC* operon, **(D)** *clpB* and, **(E)** *clpP*. Start codons are shown in green.

