## Supplementary material for "Genome-Wide Transcription Response of *Staphylococcus epidermidis* to Heat Shock and Medically Relevant Glucose Levels": Supp. Tables S1-S2

### Table S1 | *S. epidermidis* heat-shock induced transcripts.

| Functional Class & Symbol | Locus | Name | log2FC | Adjusted p-value |
|---|---|---|---|---|
| **Detoxification** | | | | |
| | F1613_RS03775 | flavocytochrome c | 3.92 | 5.04E-15 |
| | F1613_RS06270 | truncated hemoglobin YjbI | 1.90 | 1.52E-06 |
| | F1613_RS09765 | SACOL1771 family peroxiredoxin | 1.87 | 8.88E-08 |
| | F1613_RS04585 | heme-dependent peroxidase | 1.73 | 5.74E-07 |
| | F1613_RS03465 | DsrE/DsrF/DrsH-like family protein | 1.60 | 5.79E-03 |
| **DNA Integration** | | | | |
| | F1613_RS12560 | Integrase | 1.74 | 6.66E-03 |
| **DNA/RNA Repair** | | | | |
| | F1613_RS04220 | UvrB/UvrC motif-containing protein | 5.50 | 3.79E-20 |
| | F1613_RS03190 | IS1182-like element ISSep1 family transposase | 3.34 | 6.29E-13 |
| | F1613_RS05970 | IS1182-like element ISSep1 family transposase | 3.27 | 2.26E-11 |
| | F1613_RS12665 | Transposase | 3.05 | 7.94E-09 |
| dinB | F1613_RS10840 | DNA polymerase IV | 2.53 | 1.31E-11 |
| | F1613_RS03005 | recombinase family protein | 2.09 | 3.32E-04 |
| | F1613_RS11430 | single-stranded DNA-binding protein | 1.61 | 2.01E-02 |
| | F1613_RS10835 | 3'-5' exonuclease | 1.56 | 1.07E-02 |
| **Hypothetical Proteins** | | | | |
| | F1613_RS12445 | hypothetical protein | 4.77 | 1.67E-13 |
| | F1613_RS12480 | hypothetical protein | 4.17 | 1.15E-07 |
| | F1613_RS09455 | hypothetical protein | 3.59 | 1.99E-05 |
| | F1613_RS04750 | hypothetical protein | 3.34 | 5.84E-13 |
| | F1613_RS06495 | hypothetical protein | 2.67 | 1.51E-08 |
| | F1613_RS04080 | hypothetical protein | 2.40 | 6.26E-05 |
| | F1613_RS02970 | hypothetical protein | 2.39 | 1.41E-08 |
| | F1613_RS06155 | hypothetical protein | 2.37 | 1.51E-03 |
| | F1613_RS01225 | hypothetical protein | 2.34 | 4.15E-06 |
| | F1613_RS00080 | hypothetical protein | 2.32 | 9.70E-04 |
| | F1613_RS11815 | hypothetical protein | 2.32 | 1.60E-04 |
| | F1613_RS12420 | hypothetical protein | 2.32 | 4.15E-04 |
| | F1613_RS07835 | hypothetical protein | 2.24 | 1.23E-07 |
| | F1613_RS12725 | hypothetical protein | 2.18 | 2.35E-05 |
| | F1613_RS10800 | hypothetical protein | 2.12 | 9.64E-05 |
| | F1613_RS05575 | hypothetical protein | 2.06 | 9.25E-04 |
| | F1613_RS05560 | hypothetical protein | 2.05 | 1.11E-04 |
| | F1613_RS12570 | hypothetical protein | 1.98 | 1.18E-02 |
| | F1613_RS01035 | hypothetical protein | 1.92 | 1.46E-04 |
| | F1613_RS10225 | hypothetical protein | 1.86 | 7.03E-05 |
| | F1613_RS07830 | hypothetical protein | 1.77 | 3.99E-05 |
| | F1613_RS12120 | hypothetical protein | 1.66 | 6.00E-04 |
| | F1613_RS05215 | hypothetical protein | 1.63 | 1.20E-02 |
| **Metabolism** | | | | |
| lacG | F1613_RS11900 | 6-phospho-beta-galactosidase | 5.79 | 2.54E-41 |
| lacD | F1613_RS11915 | tagatose-bisphosphate aldolase | 5.79 | 1.38E-26 |
| | F1613_RS11920 | tagatose-6-phosphate kinase | 5.49 | 2.17E-27 |
| lacB | F1613_RS11925 | galactose-6-phosphate isomerase subunit LacB | 5.20 | 3.64E-25 |
| | F1613_RS03870 | ArgE/DapE family deacylase | 5.00 | 3.67E-08 |
| lacA | F1613_RS11930 | galactose-6-phosphate isomerase subunit LacA | 4.60 | 1.22E-20 |

| | | | | |
|---|---|---|---|---|
| | F1613_RS01550 | NAD(P)/FAD-dependent oxidoreductase | 4.49 | 5.54E-18 |
| | F1613_RS01545 | glutathione peroxidase | 4.37 | 9.13E-17 |
| nrdG | F1613_RS01485 | anaerobic ribonucleoside-triphosphate reductase activating protein | 4.30 | 4.32E-20 |
| nrdD | F1613_RS01490 | anaerobic ribonucleoside-triphosphate reductase | 4.23 | 4.82E-20 |
| | F1613_RS01370 | aspartate aminotransferase family protein | 4.17 | 1.38E-26 |
| ilvC | F1613_RS11225 | ketol-acid reductoisomerase | 3.82 | 5.98E-31 |
| | F1613_RS03785 | acetoin reductase | 3.72 | 2.26E-12 |
| | F1613_RS11220 | ACT domain-containing protein | 3.60 | 2.71E-10 |
| | F1613_RS04060 | thiamine pyrophosphate-dependent dehydrogenase E1 component subunit alpha | 3.55 | 0.003519281 |
| | F1613_RS04065 | alpha-ketoacid dehydrogenase subunit beta | 3.50 | 0.000297743 |
| | F1613_RS01135 | zinc-dependent alcohol dehydrogenase family protein | 3.28 | 1.02E-16 |
| | F1613_RS09310 | LLM class flavin-dependent oxidoreductase | 3.26 | 1.04E-23 |
| | F1613_RS11230 | 2-isopropylmalate synthase | 3.26 | 3.36E-19 |
| | F1613_RS06075 | NADH-dependent flavin oxidoreductase | 3.12 | 1.61E-16 |
| leuB | F1613_RS11235 | 3-isopropylmalate dehydrogenase | 3.12 | 7.41E-13 |
| pfkB | F1613_RS05155 | 1-phosphofructokinase | 3.07 | 2.21E-05 |
| gntK | F1613_RS00865 | Gluconokinase | 3.02 | 2.77E-14 |
| | F1613_RS08045 | aminodeoxychorismate/anthranilate synthase component II | 2.96 | 2.55E-06 |
| leuC | F1613_RS11240 | 3-isopropylmalate dehydratase large subunit | 2.84 | 4.28E-11 |
| pflA | F1613_RS03865 | pyruvate formate-lyase-activating protein | 2.78 | 6.97E-07 |
| | F1613_RS04620 | FAD-containing oxidoreductase | 2.76 | 1.41E-08 |
| | F1613_RS09950 | proline dehydrogenase | 2.68 | 0.000151127 |
| ilvB | F1613_RS11215 | biosynthetic-type acetolactate synthase large subunit | 2.66 | 9.89E-12 |
| pruA | F1613_RS01215 | L-glutamate gamma-semialdehyde dehydrogenase | 2.66 | 2.18E-05 |
| adhE | F1613_RS05320 | bifunctional acetaldehyde-CoA/alcohol dehydrogenase | 2.62 | 9.24E-07 |
| alsS | F1613_RS01350 | acetolactate synthase AlsS | 2.61 | 9.89E-12 |
| | F1613_RS11890 | LLM class flavin-dependent oxidoreductase | 2.52 | 1.32E-12 |
| | F1613_RS04070 | dihydrolipoamide acetyltransferase family protein | 2.49 | 2.13E-03 |
| | F1613_RS01000 | aldehyde dehydrogenase family protein | 2.49 | 8.05E-04 |
| | F1613_RS01355 | L-lactate dehydrogenase | 2.43 | 4.87E-11 |
| leuD | F1613_RS11245 | 3-isopropylmalate dehydratase small subunit | 2.40 | 1.34E-09 |
| | F1613_RS02610 | PLP-dependent aspartate aminotransferase family protein | 2.39 | 4.74E-09 |
| | F1613_RS06705 | oleate hydratase | 2.38 | 5.16E-03 |
| budA | F1613_RS01345 | acetolactate decarboxylase | 2.35 | 7.57E-10 |
| ilvA | F1613_RS11250 | threonine ammonia-lyase IlvA | 2.35 | 1.37E-14 |
| | F1613_RS00685 | 2-dehydropantoate 2-reductase | 2.35 | 2.89E-11 |
| dhaK | F1613_RS03965 | dihydroxyacetone kinase subunit DhaK | 2.32 | 7.57E-09 |
| | F1613_RS03770 | FAD:protein FMN transferase | 2.29 | 6.60E-08 |
| pflB | F1613_RS03860 | formate C-acetyltransferase | 2.23 | 1.90E-05 |
| dhaL | F1613_RS03970 | dihydroxyacetone kinase subunit DhaL | 2.17 | 3.24E-07 |
| zwf | F1613_RS08725 | glucose-6-phosphate dehydrogenase | 2.14 | 1.27E-08 |
| trpD | F1613_RS08050 | anthranilate phosphoribosyltransferase | 2.10 | 1.12E-06 |
| | F1613_RS04185 | arylamine N-acetyltransferase | 2.08 | 8.10E-06 |
| ald | F1613_RS09695 | alanine dehydrogenase | 2.07 | 6.95E-06 |
| | F1613_RS01745 | (S)-acetoin forming diacetyl reductase | 2.07 | 7.75E-05 |
| dhaM | F1613_RS03975 | dihydroxyacetone kinase phosphoryl donor subunit DhaM | 2.04 | 3.29E-06 |
| | F1613_RS03960 | glycerol dehydrogenase | 2.01 | 3.88E-06 |
| bshC | F1613_RS07090 | bacillithiol biosynthesis cysteine-adding enzyme BshC | 1.96 | 1.39E-08 |
| | F1613_RS08740 | alpha-glucosidase | 1.95 | 4.58E-07 |
| | F1613_RS09305 | cysteine desulfurase family protein | 1.94 | 2.52E-06 |
| | F1613_RS02605 | PLP-dependent transferase | 1.94 | 8.26E-08 |
| | F1613_RS02260 | glutamate synthase subunit beta | 1.92 | 8.97E-05 |

| | | | | |
|---|---|---|---|---|
| | F1613_RS05795 | thioredoxin family protein | 1.91 | 3.23E-05 |
| | F1613_RS01010 | NAD(P)H-dependent oxidoreductase | 1.91 | 4.30E-09 |
| arcC | F1613_RS03315 | carbamate kinase | 1.90 | 1.07E-03 |
| | F1613_RS08040 | anthranilate synthase component I | 1.87 | 1.77E-05 |
| | F1613_RS03370 | aspartate aminotransferase family protein | 1.85 | 3.49E-10 |
| | F1613_RS01670 | alpha-keto acid decarboxylase family protein | 1.80 | 1.06E-03 |
| | F1613_RS00940 | thiazole synthase | 1.75 | 1.22E-04 |
| | F1613_RS00620 | 2,3-diphosphoglycerate-dependent phosphoglycerate mutase | 1.74 | 1.04E-02 |
| argH | F1613_RS06100 | argininosuccinate lyase | 1.73 | 2.97E-03 |
| | F1613_RS08195 | alanine racemase | 1.72 | 8.59E-03 |
| | F1613_RS01635 | poly-gamma-glutamate hydrolase family protein | 1.71 | 1.94E-06 |
| thrB | F1613_RS07855 | homoserine kinase | 1.69 | 2.57E-08 |
| betA | F1613_RS01460 | betaine-aldehyde dehydrogenase | 1.68 | 8.73E-03 |
| | F1613_RS07840 | aspartate kinase | 1.68 | 1.12E-03 |
| gpmI | F1613_RS05600 | 2,3-bisphosphoglycerate-independent phosphoglycerate mutase | 1.67 | 4.34E-04 |
| thiF | F1613_RS00945 | thiazole biosynthesis adenylyltransferase ThiF | 1.67 | 1.94E-05 |
| thiO | F1613_RS00930 | glycine oxidase ThiO | 1.63 | 2.60E-04 |
| thiS | F1613_RS00935 | sulfur carrier protein ThiS | 1.62 | 2.10E-03 |
| | F1613_RS02615 | bifunctional homocysteine S-methyltransferase/methylenetetrahydrofolate reductase | 1.62 | 9.07E-05 |
| ilvD | F1613_RS11210 | dihydroxy-acid dehydratase | 1.61 | 4.55E-06 |
| tpiA | F1613_RS05595 | triose-phosphate isomerase | 1.61 | 1.74E-04 |
| nagB | F1613_RS04495 | glucosamine-6-phosphate deaminase | 1.59 | 1.53E-05 |
| | F1613_RS05590 | phosphoglycerate kinase | 1.58 | 3.64E-04 |
| | F1613_RS08710 | SDR family NAD(P)-dependent oxidoreductase | 1.58 | 7.92E-04 |
| adhP | F1613_RS04670 | alcohol dehydrogenase AdhP | 1.57 | 2.83E-03 |
| pcp | F1613_RS04095 | pyroglutamyl-peptidase I | 1.54 | 4.71E-05 |
| | F1613_RS00035 | 2-hydroxyacid dehydrogenase family protein | 1.52 | 4.32E-04 |
| **Molecular Chaperones** | | | | |
| clpB | F1613_RS06180 | ATP-dependent chaperone ClpB | 6.15 | 6.64E-17 |
| | F1613_RS04230 | ATP-dependent Clp protease ATP-binding subunit clpC | 5.73 | 4.24E-20 |
| dnaK | F1613_RS09115 | molecular chaperone DnaK | 3.91 | 5.29E-15 |
| grpE | F1613_RS09120 | nucleotide exchange factor GrpE | 3.77 | 8.23E-14 |
| groL | F1613_RS11085 | chaperonin GroEL | 3.51 | 3.99E-17 |
| groES | F1613_RS11090 | co-chaperone GroES | 3.37 | 6.96E-16 |
| dnaJ | F1613_RS09110 | molecular chaperone DnaJ | 2.97 | 2.97E-16 |
| hslO | F1613_RS02040 | Hsp33 family molecular chaperone HslO | 2.05 | 1.68E-04 |
| **Ribosome Biogenesis** | | | | |
| prmA | F1613_RS09105 | 50S ribosomal protein L11 methyltransferase | 2.95 | 4.32E-14 |
| **Oxidative Phosphorylation** | | | | |
| | F1613_RS06745 | cytochrome ubiquinol oxidase subunit I | 4.99 | 2.62E-31 |
| | F1613_RS06750 | cytochrome d ubiquinol oxidase subunit II | 4.97 | 4.11E-28 |
| **Protein Degradation** | | | | |
| | F1613_RS04225 | protein arginine kinase | 5.63 | 9.87E-20 |
| clpP | F1613_RS05555 | ATP-dependent Clp endopeptidase proteolytic subunit ClpP | 3.16 | 1.90E-17 |
| mecA | F1613_RS06250 | adaptor protein MecA | 1.94 | 1.32E-03 |
| yjbH | F1613_RS06265 | protease adaptor protein YjbH | 1.78 | 6.45E-08 |
| | F1613_RS08220 | CPBP family intramembrane metalloprotease | 1.76 | 1.62E-02 |
| | F1613_RS09795 | S1C family serine protease | 1.61 | 2.10E-03 |

| | | | | |
|---|---|---|---|---|
| **Regulation** | | | | |
| | F1613_RS04215 | CtsR family transcription regulator | 5.38 | 5.02E-22 |
| | F1613_RS01555 | MarR family transcription regulator | 4.90 | 1.29E-23 |
| hrcA | F1613_RS09125 | heat-inducible transcriptional repressor HrcA | 3.83 | 1.35E-15 |
| | F1613_RS04625 | Rrf2 family transcription regulator | 2.94 | 9.70E-08 |
| | F1613_RS03175 | metalloregulator ArsR/SmtB family transcription factor | 2.67 | 2.72E-05 |
| | F1613_RS11940 | NAD-dependent protein deacylase | 2.66 | 2.56E-15 |
| | F1613_RS01755 | BglG family transcription antiterminator | 2.38 | 2.92E-10 |
| | F1613_RS00870 | GntR family transcription regulator | 2.37 | 2.86E-07 |
| | F1613_RS05150 | DeoR/GlpR family DNA-binding transcription regulator | 2.33 | 7.16E-05 |
| saeR | F1613_RS05195 | response regulator transcription factor SaeR | 2.27 | 1.06E-03 |
| raiA | F1613_RS05465 | ribosome-associated translation inhibitor RaiA | 2.26 | 3.45E-05 |
| vraR | F1613_RS10785 | two-component system response regulator VraR | 2.20 | 4.58E-07 |
| | F1613_RS10000 | competence protein ComK | 2.20 | 1.25E-04 |
| | F1613_RS09995 | sigma-70 family RNA polymerase sigma factor | 2.17 | 3.48E-04 |
| | F1613_RS06475 | competence protein ComK | 2.17 | 2.29E-04 |
| | F1613_RS00070 | HTH domain-containing protein | 2.06 | 1.02E-06 |
| | F1613_RS02270 | LysR family transcription regulator | 1.99 | 6.00E-07 |
| | F1613_RS03505 | metalloregulator ArsR/SmtB family transcription factor | 1.79 | 8.05E-05 |
| | F1613_RS03130 | metalloregulator ArsR/SmtB family transcription factor | 1.61 | 1.52E-03 |
| | F1613_RS12175 | winged helix DNA-binding protein | 1.57 | 1.66E-02 |
| | F1613_RS02675 | GNAT family protein | 1.55 | 1.19E-04 |
| **Secretion** | | | | |
| | F1613_RS10230 | TrbC/VirB2 family protein | 2.16 | 5.81E-09 |
| asp3 | F1613_RS01845 | accessory Sec system protein Asp3 | 1.75 | 2.19E-07 |
| secA2 | F1613_RS01840 | accessory Sec system translocase SecA2 | 1.52 | 3.51E-06 |
| **Signal Transduction** | | | | |
| | F1613_RS05190 | HAMP domain-containing sensor histidine kinase | 2.37 | 8.89E-04 |
| | F1613_RS10790 | sensor histidine kinase | 2.00 | 1.48E-08 |
| **Stress Response** | | | | |
| liaF | F1613_RS10795 | cell wall-active antibiotics response protein LiaF | 1.75 | 9.78E-06 |
| | F1613_RS09700 | universal stress protein | 1.70 | 2.49E-05 |
| | F1613_RS09680 | universal stress protein | 1.55 | 1.29E-03 |
| **Surface Protein** | | | | |
| | F1613_RS06965 | YSIRK-type signal peptide-containing protein | 1.86 | 1.93E-05 |
| **Translation** | | | | |
| | F1613_RS10325 | tRNA-Gly | 1.92 | 1.63E-03 |
| | F1613_RS10535 | tRNA-Cys | 1.51 | 1.64E-02 |
| | F1613_RS10330 | tRNA-His | 1.50 | 2.63E-02 |
| **Transport** | | | | |
| | F1613_RS11905 | lactose-specific PTS transporter subunit EIIC | 5.94 | 2.24E-33 |
| | F1613_RS11910 | PTS lactose/cellobiose transporter subunit IIA | 5.77 | 8.75E-29 |
| | F1613_RS03780 | MFS transporter | 4.43 | 4.74E-21 |
| | F1613_RS03185 | heavy metal translocating P-type ATPase | 3.79 | 1.50E-07 |
| | F1613_RS05160 | PTS fructose transporter subunit IIABC | 3.72 | 3.85E-06 |
| | F1613_RS06755 | TrkA family potassium uptake protein | 3.11 | 6.97E-16 |
| | F1613_RS01230 | heavy metal translocating P-type ATPase | 3.05 | 8.08E-11 |
| cydC | F1613_RS05080 | thiol reductant ABC exporter subunit CydC | 3.02 | 3.85E-15 |
| | F1613_RS05075 | ABC transporter ATP-binding protein/permease | 2.85 | 8.17E-15 |
| | F1613_RS08655 | ECF transporter S component | 2.63 | 4.33E-09 |
| | F1613_RS03925 | YfcC family protein | 2.27 | 1.12E-06 |
| | F1613_RS03435 | SLC45 family MFS transporter | 2.27 | 1.05E-10 |
| | F1613_RS04815 | sodium:proton antiporter | 2.20 | 3.02E-13 |

| | | | | |
|---|---|---|---|---|
| | F1613_RS02280 | YibE/F family protein | 2.14 | 5.84E-11 |
| | F1613_RS03180 | ZIP family metal transporter | 2.13 | 4.26E-04 |
| | F1613_RS08210 | ATP-binding cassette domain-containing protein | 1.88 | 1.35E-03 |
| | F1613_RS08205 | ABC transporter permease | 1.72 | 6.96E-03 |
| | F1613_RS10695 | ABC transporter ATP-binding protein | 1.71 | 1.73E-04 |
| | F1613_RS03910 | osmoprotectant ABC transporter substrate-binding protein | 1.70 | 1.31E-06 |
| arsA | F1613_RS03495 | arsenical pump-driving ATPase | 1.68 | 7.17E-05 |
| | F1613_RS03915 | ABC transporter permease | 1.68 | 4.39E-06 |
| yut | F1613_RS12295 | urea transporter | 1.64 | 1.23E-03 |
| arsD | F1613_RS03500 | arsenite efflux transporter metallochaperone ArsD | 1.62 | 3.60E-04 |
| rarD | F1613_RS04040 | EamA family transporter RarD | 1.61 | 2.10E-03 |
| | F1613_RS02275 | YibE/F family protein | 1.56 | 3.52E-03 |
| | F1613_RS10095 | Ltp family lipoprotein | 1.56 | 1.53E-03 |
| | F1613_RS05000 | globin domain-containing protein | 1.52 | 2.11E-03 |
| | F1613_RS04755 | DMT family transporter | 1.52 | 4.03E-04 |
| | F1613_RS06820 | ABC transporter permease | 1.52 | 2.48E-04 |
| | F1613_RS00665 | amino acid permease | 1.51 | 3.13E-05 |
| | F1613_RS04030 | ABC-ATPase domain-containing protein | 1.51 | 4.08E-05 |
| **tRNA Biosynthesis** | | | | |
| mnmA | F1613_RS09300 | tRNA 2-thiouridine(34) synthase MnmA | 1.54 | 1.89E-03 |
| **Unknown Function** | | | | |
| | F1613_RS04110 | YceI family protein | 4.01 | 1.32E-19 |
| | F1613_RS12660 | uncharacterized gene | 3.91 | 6.02E-21 |
| | F1613_RS11080 | cell wall surface anchor protein | 3.79 | 3.50E-17 |
| | F1613_RS07270 | TM2 domain-containing protein | 3.19 | 3.85E-06 |
| | F1613_RS08215 | Msa family membrane protein | 3.06 | 0.000107032 |
| | F1613_RS02870 | alpha/beta fold hydrolase | 2.66 | 3.51E-05 |
| | F1613_RS12185 | uncharacterized gene | 2.62 | 0.001381984 |
| | F1613_RS12630 | uncharacterized gene | 2.37 | 3.88E-06 |
| | F1613_RS12510 | uncharacterized gene | 2.35 | 1.36E-05 |
| | F1613_RS05200 | DoxX family protein | 2.28 | 1.39E-03 |
| | F1613_RS04970 | Bax inhibitor-1 family protein | 2.27 | 1.19E-07 |
| | F1613_RS05565 | TIGR01777 family oxidoreductase | 2.20 | 1.27E-08 |
| | F1613_RS07920 | DUF896 domain-containing protein | 2.18 | 9.44E-08 |
| | F1613_RS05205 | DM13 domain-containing protein | 2.17 | 2.97E-03 |
| | F1613_RS03145 | uncharacterized gene | 1.99 | 2.82E-03 |
| | F1613_RS12190 | uncharacterized gene | 1.95 | 1.66E-05 |
| | F1613_RS09745 | GAF domain-containing protein | 1.78 | 1.96E-06 |
| | F1613_RS12520 | uncharacterized gene | 1.77 | 1.74E-03 |
| | F1613_RS01120 | alpha/beta hydrolase | 1.66 | 3.89E-04 |
| yidD | F1613_RS10075 | membrane protein insertion efficiency factor YidD | 1.61 | 1.90E-04 |
| | F1613_RS06170 | metal-sulfur cluster assembly factor | 1.58 | 4.71E-05 |
| | F1613_RS03920 | alpha/beta hydrolase family protein | 1.54 | 4.87E-05 |
| | F1613_RS03440 | uncharacterized gene | 1.54 | 1.55E-03 |
| | F1613_RS12555 | uncharacterized gene | 1.51 | 3.13E-04 |
| **Urease Accessory Proteins** | | | | |
| | F1613_RS12320 | urease accessory protein UreF | 1.59 | 3.99E-03 |
| ureE | F1613_RS12315 | urease accessory protein UreE | 1.52 | 5.22E-03 |
| | F1613_RS12330 | urease accessory protein UreD | 1.50 | 4.91E-03 |
| **Virulence Factors** | | | | |
| vraX | F1613_RS04535 | C1q-binding complement inhibitor VraX | 5.42 | 3.72E-10 |

| | | | | |
|---|---|---|---|---|
| blaZ | F1613_RS10980 | penicillin-hydrolyzing class A beta-lactamase BlaZ | 4.19 | 9.08E-13 |
| blaI | F1613_RS10970 | penicillinase repressor BlaI | 2.32 | 5.68E-08 |
| blaR1 | F1613_RS10975 | beta-lactam sensor/signal transducer BlaR1 | 2.25 | 3.96E-06 |

### Table S2| *S. epidermidis* heat-shock repressed transcripts.

| Functional Class & Symbol | Locus | Name | log2FC | Adjusted p-value |
|---|---|---|---|---|
| **Beta-Class Phenol-Soluble Modulin** | | | | |
| | F1613_RS07075 | beta-class phenol-soluble modulin | -2.20 | 2.87E-03 |
| | F1613_RS07060 | beta-class phenol-soluble modulin | -1.89 | 8.67E-03 |
| | F1613_RS07065 | beta-class phenol-soluble modulin | -1.63 | 1.90E-02 |
| | F1613_RS07070 | beta-class phenol-soluble modulin | -1.55 | 2.47E-02 |
| **Cell Wall Structure** | | | | |
| | F1613_RS05940 | teichoic acid D-Ala incorporation-associated protein DltX | -4.40 | 1.27E-08 |
| dltA | F1613_RS05945 | D-alanine--poly(phosphoribitol) ligase subunit DltA | -3.73 | 1.04E-05 |
| dltB | F1613_RS05950 | D-alanyl-lipoteichoic acid biosynthesis protein DltB | -3.63 | 2.18E-05 |
| dltC | F1613_RS05955 | D-alanine--poly(phosphoribitol) ligase subunit 2 | -3.38 | 5.78E-05 |
| dltD | F1613_RS05960 | D-alanyl-lipoteichoic acid biosynthesis protein DltD | -3.34 | 6.11E-06 |
| | F1613_RS01265 | transglycosylase family protein | -3.08 | 2.98E-05 |
| **DNA Restriction-Modification System** | | | | |
| | F1613_RS03040 | restriction endonuclease subunit R | -1.53 | 3.87E-02 |
| **DNA/RNA Repair** | | | | |
| ssb | F1613_RS02565 | single-stranded DNA-binding protein | -3.06 | 1.81E-11 |
| | F1613_RS09000 | deoxyribonuclease IV | -1.66 | 6.76E-03 |
| radC | F1613_RS09460 | DNA repair protein RadC | -1.59 | 4.52E-03 |
| **Hypothetical Protein** | | | | |
| | F1613_RS00595 | hypothetical protein | -2.90 | 7.05E-13 |
| | F1613_RS10130 | hypothetical protein | -2.60 | 2.50E-03 |
| | F1613_RS03300 | hypothetical protein | -2.55 | 2.06E-09 |
| | F1613_RS05655 | hypothetical protein | -2.27 | 7.58E-12 |
| | F1613_RS07970 | hypothetical protein | -2.23 | 3.40E-06 |
| | F1613_RS01415 | hypothetical protein | -2.14 | 2.15E-06 |
| | F1613_RS04530 | hypothetical protein | -1.98 | 1.85E-03 |
| | F1613_RS05640 | hypothetical protein | -1.95 | 2.57E-07 |
| | F1613_RS10300 | hypothetical protein | -1.82 | 2.69E-02 |
| | F1613_RS10270 | hypothetical protein | -1.80 | 6.56E-03 |
| | F1613_RS05665 | hypothetical protein | -1.79 | 2.48E-03 |
| | F1613_RS02945 | hypothetical protein | -1.78 | 5.50E-06 |
| | F1613_RS12485 | hypothetical protein | -1.77 | 6.93E-05 |
| | F1613_RS12415 | hypothetical protein | -1.71 | 6.67E-10 |
| | F1613_RS00295 | hypothetical protein | -1.63 | 1.55E-03 |
| | F1613_RS11670 | hypothetical protein | -1.58 | 1.08E-06 |
| **Metabolism** | | | | |
| | F1613_RS03290 | NADP-dependent oxidoreductase | -3.35 | 2.30E-15 |
| | F1613_RS10115 | Amidase | -2.80 | 7.12E-05 |
| | F1613_RS11355 | DEAD/DEAH box helicase | -2.74 | 2.11E-06 |
| | F1613_RS07205 | aspartate carbamoyltransferase catalytic subunit | -2.56 | 4.68E-08 |
| | F1613_RS00215 | galactose mutarotase | -2.40 | 2.29E-10 |
| | F1613_RS01150 | pyruvate oxidase | -2.18 | 5.95E-04 |
| plsY | F1613_RS07975 | glycerol-3-phosphate 1-O-acyltransferase PlsY | -2.17 | 1.83E-10 |

| | | | | |
|---|---|---|---|---|
| | F1613_RS04320 | class I SAM-dependent methyltransferase | -2.13 | 5.08E-04 |
| | F1613_RS04425 | deoxynucleoside kinase | -2.00 | 1.50E-05 |
| | F1613_RS04420 | deoxynucleoside kinase | -1.99 | 2.73E-05 |
| | F1613_RS11585 | CTP synthase | -1.97 | 6.08E-11 |
| | F1613_RS07080 | YjjG family noncanonical pyrimidine nucleotidase | -1.93 | 6.33E-05 |
| rpe | F1613_RS07310 | ribulose-phosphate 3-epimerase | -1.84 | 5.08E-04 |
| | F1613_RS09005 | DEAD/DEAH box helicase | -1.82 | 3.48E-03 |
| | F1613_RS07500 | isoprenyl transferase | -1.57 | 4.14E-04 |
| | F1613_RS04520 | acetyl-CoA C-acyltransferase | -1.57 | 2.21E-05 |
| galU | F1613_RS00855 | UTP--glucose-1-phosphate uridylyltransferase GalU | -1.56 | 7.96E-05 |
| rnmV | F1613_RS02140 | ribonuclease M5 | -1.54 | 8.77E-03 |
| coaW | F1613_RS11600 | type II pantothenate kinase | -1.54 | 8.24E-06 |
| | F1613_RS07215 | carbamoyl phosphate synthase small subunit | -1.51 | 6.19E-05 |
| **Metal-Binding Proteins** | | | | |
| | F1613_RS09010 | Nif3-like dinuclear metal center hexameric protein | -1.81 | 3.82E-04 |
| **DNA Metabolic Process** | | | | |
| | F1613_RS05645 | tyrosine-type recombinase/integrase | -2.69 | 2.39E-05 |
| **Regulation** | | | | |
| | F1613_RS10440 | helix-turn-helix transcription regulator | -3.08 | 1.57E-12 |
| rsp | F1613_RS00450 | AraC family transcription regulator Rsp | -2.47 | 2.96E-05 |
| | F1613_RS11065 | GntR family transcription regulator | -2.20 | 5.03E-04 |
| pyrR | F1613_RS07195 | bifunctional pyr operon transcription regulator/uracil phosphoribosyltransferase PyrR | -2.04 | 5.00E-03 |
| | F1613_RS09035 | helix-turn-helix transcription regulator | -1.81 | 1.58E-03 |
| **Ribosome Biogenesis** | | | | |
| rsgA | F1613_RS07305 | ribosome small subunit-dependent GTPase A | -1.73 | 3.23E-07 |
| **Stress Response** | | | | |
| | F1613_RS05710 | cold-shock protein | -3.18 | 1.16E-08 |
| cspA | F1613_RS08225 | cold shock protein CspA | -2.71 | 2.21E-05 |
| | F1613_RS02545 | GlsB/YeaQ/YmgE family stress response membrane protein | -1.96 | 1.30E-03 |
| **Transcription Machinery** | | | | |
| rpoE | F1613_RS11590 | DNA-directed RNA polymerase subunit delta | -1.94 | 8.91E-07 |
| **Ribosome/ Translation** | | | | |
| rpsF | F1613_RS02570 | 30S ribosomal protein S6 | -3.05 | 2.58E-10 |
| | F1613_RS01980 | *tRNA-Arg | -3.01 | 3.86E-07 |
| rpsR | F1613_RS02560 | 30S ribosomal protein S18 | -2.82 | 9.62E-06 |
| | F1613_RS11735 | tRNA-Lys | -2.62 | 1.61E-04 |
| typA | F1613_RS06860 | translational GTPase TypA | -2.49 | 1.08E-06 |
| | F1613_RS01985 | *tRNA-Leu | -2.47 | 1.28E-05 |
| infC | F1613_RS09555 | translation initiation factor IF-3 | -2.27 | 2.13E-08 |
| rpmI | F1613_RS09550 | 50S ribosomal protein L35 | -2.26 | 8.10E-06 |
| rplJ | F1613_RS04310 | 50S ribosomal protein L10 | -2.24 | 1.64E-04 |
| | F1613_RS09040 | glycine--tRNA ligase | -2.18 | 9.70E-05 |
| rplT | F1613_RS09545 | 50S ribosomal protein L20 | -2.17 | 4.26E-04 |
| rpsO | F1613_RS07565 | 30S ribosomal protein S15 | -2.16 | 1.26E-04 |
| rplL | F1613_RS04315 | 50S ribosomal protein L7/L12 | -2.02 | 1.31E-03 |
| | F1613_RS10455 | RluA family pseudouridine synthase | -2.02 | 1.61E-08 |

| | | | | |
|---|---|---|---|---|
| ileS | F1613_RS07170 | isoleucine--tRNA ligase | -1.76 | 2.15E-05 |
| thrS | F1613_RS09565 | threonine--tRNA ligase | -1.65 | 6.30E-03 |
| serS | F1613_RS02850 | serine--tRNA ligase | -1.51 | 1.15E-03 |
| **Transport** | | | | |
| | F1613_RS03285 | ABC transporter permease | -3.47 | 6.35E-17 |
| | F1613_RS03280 | ABC transporter ATP-binding protein | -3.33 | 7.19E-13 |
| | F1613_RS07200 | NCS2 family nucleobase:cation symporter | -3.23 | 5.01E-05 |
| | F1613_RS00310 | HlyD family efflux transporter periplasmic adaptor subunit | -2.87 | 6.45E-08 |
| ptsG | F1613_RS01145 | glucose-specific PTS transporter subunit IIBC | -2.77 | 1.44E-06 |
| | F1613_RS00745 | amino acid permease | -2.55 | 2.83E-06 |
| | F1613_RS00305 | DHA2 family efflux MFS transporter permease subunit | -2.46 | 5.64E-06 |
| | F1613_RS05370 | ABC transporter permease | -2.22 | 8.18E-04 |
| | F1613_RS08160 | ATP-binding cassette domain-containing protein | -2.22 | 4.12E-08 |
| pmtA | F1613_RS11060 | phenol-soluble modulin export ABC transporter ATP-binding protein PmtA | -2.12 | 4.32E-04 |
| | F1613_RS03990 | MDR family MFS transporter | -2.06 | 4.17E-05 |
| | F1613_RS05375 | iron chelate uptake ABC transporter family permease subunit | -2.03 | 5.52E-04 |
| | F1613_RS05380 | ATP-binding cassette domain-containing protein | -2.02 | 4.98E-04 |
| | F1613_RS10505 | PTS transporter subunit IIC | -2.01 | 1.25E-10 |
| pmtB | F1613_RS11055 | phenol-soluble modulin export ABC transporter permease subunit PmtB | -2.01 | 6.48E-04 |
| | F1613_RS05385 | siderophore ABC transporter substrate-binding protein | -1.90 | 8.92E-04 |
| | F1613_RS11970 | energy-coupling factor transporter ATPase | -1.81 | 5.01E-05 |
| | F1613_RS11840 | Fe(3+) dicitrate ABC transporter substrate-binding protein | -1.80 | 6.41E-03 |
| | F1613_RS09635 | amino acid permease | -1.78 | 1.83E-05 |
| | F1613_RS04525 | protein VraC | -1.76 | 3.81E-03 |
| pmtC | F1613_RS11050 | phenol-soluble modulin export ABC transporter ATP-binding protein PmtC | -1.68 | 5.58E-03 |
| | F1613_RS02750 | RND transporter | -1.65 | 1.37E-02 |
| | F1613_RS00160 | Na+/H+ antiporter NhaC family protein | -1.54 | 9.05E-06 |
| mscL | F1613_RS07950 | large conductance mechanosensitive channel protein MscL | -1.55 | 6.45E-08 |
| **tRNA Biosynthesis** | | | | |
| | F1613_RS09015 | tRNA (adenine(22)-N(1))-methyltransferase TrmK | -1.94 | 5.14E-05 |
| rsmA | F1613_RS02135 | 16S rRNA (adenine(1518)-N(6)/adenine(1519)-N(6))-dimethyltransferase RsmA | -1.69 | 6.43E-04 |
| | F1613_RS01310 | epoxyqueuosine reductase QueH | -1.62 | 3.53E-03 |
| | F1613_RS02165 | tRNA1(Val) (adenine(37)-N6)-methyltransferase | -1.56 | 3.51E-04 |
| **Unknown Function** | | | | |
| | F1613_RS03295 | HXXEE domain-containing protein | -3.20 | 3.60E-16 |
| | F1613_RS10435 | DUF445 family protein | -2.78 | 5.43E-09 |
| | F1613_RS10135 | AAA family ATPase | -2.78 | 6.32E-04 |
| | F1613_RS02640 | sterile alpha motif-like domain-containing protein | -2.36 | 3.52E-03 |
| | F1613_RS11185 | AAA family ATPase | -2.20 | 1.19E-04 |
| | F1613_RS01255 | CHAP domain-containing protein | -2.16 | 1.81E-03 |
| | F1613_RS01155 | LrgB family protein | -2.14 | 2.61E-04 |
| | F1613_RS05980 | YuzD family protein | -2.11 | 5.16E-03 |
| | F1613_RS02400 | VOC family protein | -2.03 | 4.23E-04 |
| | F1613_RS04965 | LysM peptidoglycan-binding domain-containing protein | -2.02 | 5.53E-05 |
| | F1613_RS09160 | helix-hairpin-helix domain-containing protein | -1.99 | 1.06E-11 |
| | F1613_RS12355 | CHAP domain-containing protein | -1.95 | 5.56E-03 |
| | F1613_RS05400 | GrpB family protein | -1.94 | 8.77E-03 |
| | F1613_RS03270 | HXXEE domain-containing protein | -1.88 | 7.30E-04 |

| | | | | |
|---|---|---|---|---|
| | F1613_RS04510 | HAD family hydrolase | -1.86 | 1.34E-02 |
| | F1613_RS00015 | DUF4870 domain-containing protein | -1.83 | 2.60E-08 |
| | F1613_RS00590 | Aminoacyltransferase | -1.81 | 7.80E-07 |
| | F1613_RS04515 | uncharacterized gene | -1.78 | 4.23E-04 |
| | F1613_RS06565 | acyltransferase family protein | -1.75 | 7.43E-08 |
| | F1613_RS07820 | membrane protein | -1.60 | 5.04E-03 |
| | F1613_RS12655 | uncharacterized gene | -1.56 | 2.90E-02 |
| | F1613_RS01730 | DUF1672 domain-containing protein | -1.52 | 5.96E-03 |
| | F1613_RS04915 | membrane protein | -1.51 | 3.01E-09 |
| **Virulence Factors** | | | | |
| mprF | F1613_RS08005 | bifunctional lysylphosphatidylglycerol flippase/synthetase MprF | -1.74 | 3.71E-03 |
